# The Pex6 N1 domain is required for Pex15 binding and proper assembly with Pex1

**DOI:** 10.1101/2023.09.15.557798

**Authors:** Bashir A. Ali, Ryan M. Judy, Saikat Chowdhury, Nicole K. Jacobsen, Dominic T. Castanzo, Kaili L. Carr, Chris D. Richardson, Gabriel C. Lander, Andreas Martin, Brooke M. Gardner

## Abstract

The heterohexameric AAA-ATPase Pex1/Pex6 is essential for the formation and maintenance of peroxisomes. Pex1/Pex6, similar to other AAA-ATPases, uses the energy from ATP hydrolysis to mechanically thread substrate proteins through its central pore, thereby unfolding them. In related AAA-ATPase motors, substrates are recruited through binding to the motor’s N-terminal domains or N-terminally bound co-factors. Here we use structural and biochemical techniques to characterize the function of the N1 domain in Pex6 from budding yeast, *S. cerevisiae*. We found that although Pex1/ΔN1-Pex6 is an active ATPase *in vitro*, it does not support Pex1/Pex6 function at the peroxisome *in vivo*. An X-ray crystal structure of the isolated Pex6 N1 domain shows that the Pex6 N1 domain shares the same fold as the N terminal domains of PEX1, CDC48, or NSF, despite poor sequence conservation. Integrating this structure with a cryo-EM reconstruction of Pex1/Pex6, AlphaFold2 predictions, and biochemical assays shows that Pex6 N1 mediates binding to both the peroxisomal membrane tether Pex15 and an extended loop from the D2 ATPase domain of Pex1 that influences Pex1/Pex6 heterohexamer stability. Given the direct interactions with both Pex15 and the D2 ATPase domains, the Pex6 N1 domain is poised to coordinate binding of co-factors and substrates with Pex1/Pex6 ATPase activity.

## Introduction

Peroxisomes are membrane-bound organelles that harbor enzymes for specialized metabolic reactions, such as the β-oxidation of very long chain fatty acids. The biogenesis and maintenance of peroxisomes depends on approximately 35 Pex proteins [Smith and Aitchison 2013], including Pex1 and Pex6. Pex1 and Pex6 belong to a family of Type 2 AAA-ATPases that form hexameric complexes consisting of two stacked ATPase rings, D1 and D2, topped by N-terminal domains. These AAA-ATPase motor proteins translate the chemical energy of nucleotide hydrolysis to exert mechanical force on their substrates, as typified by Cdc48’s extraction of proteins from the endoplasmic reticulum (ER) in ER-associated degradation and NSF’s disassembly of SNARE complexes after vesicle fusion. Unlike these better-studied homohexameric AAA-ATPases, Pex1 and Pex6 form a heterohexameric ATPase with alternating Pex1 and Pex6 subunits [Gardner et al 2015, Blok et al 2015, Ciniawsky et al 2015]. This heterohexameric assembly creates two unique interfaces: the Pex1:Pex6 interface, which contains the nucleotide binding sites including the Walker A and Walker B motifs provided by the large AAA subdomains of Pex1, while Pex6 contributes the arginine finger motif; and the Pex6:Pex1 interface, which contains the nucleotide binding sites, Walker A, and Walker B motifs provided by the large AAA subdomain of Pex6, while Pex1 contributes the arginine finger motif. Additionally, Pex1 and Pex6 each have two N-terminal domains (N1 and N2, **Figure 1A**) [Blok et al 2015] which is in contrast to Cdc48 and NSF with only a single N-terminal domain per protomer. Thus, this duplication and the heterohexameric assembly increases the number of N-terminal domains available to recruit substrates and co-factors to the motor, potentially allowing for specialization. Characterizing the N-terminal domains of Pex1/Pex6 therefore has the potential to reveal novel or unanticipated functions of the Pex1/Pex6 heterohexamer, and thus we set out to elucidate the role of the Pex6 N1 domain in Pex1/Pex6 motor function.

**Figure 1.**
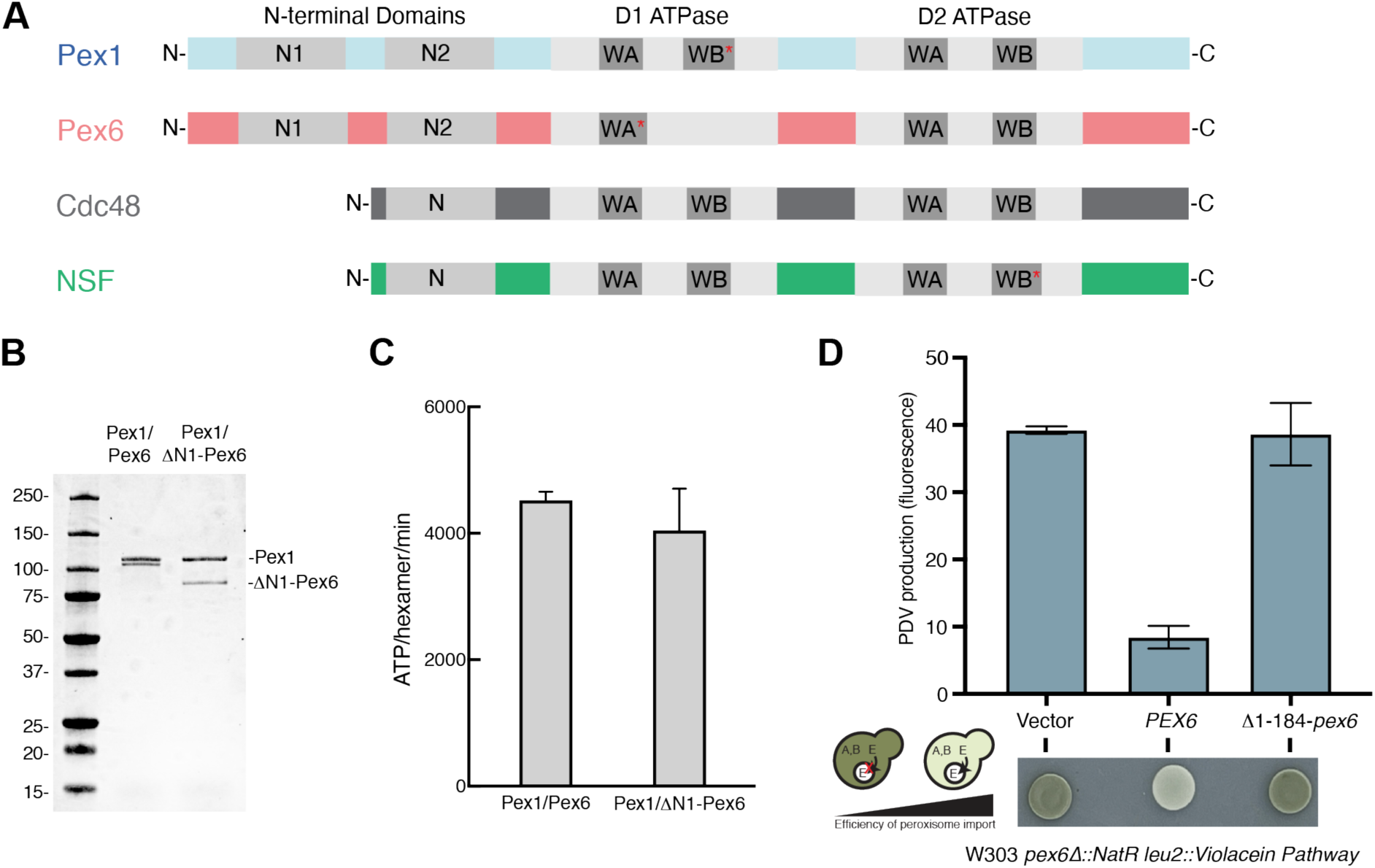
A) Domain architecture of Pex1, Pex6, Cdc48, and NSF showing N-terminal substrate/cofactor binding domains and ATPase domains, D1 and D2. Red asterisks indicate endogenous substitutions that disrupt ATPase activity. B) SDS-PAGE analysis of tandem-affinity purified Pex1/Pex6 and Pex1/ΔN1-Pex6. C) ATPase activity of Pex1/Pex6 and Pex1/ΔN1-Pex6. Shown are the average and standard deviation of three technical replicates. D) The sequential activity of VioA, VioB, and VioE produce prodeoxyviolacein (PDV), a green pigment that can be visualized by colony color on a plate and fluorescence in yeast extract. When VioE is targeted to the peroxisome by a C-terminal peroxisome targeting signal, the amount of PDV is inversely correlated with the efficiency of peroxisome import. Complementation of *Δpex6* with plasmids expressing *PEX6* shows that *ΔN1-PEX6* causes a peroxisome import defect, with high PDV production. Average and standard deviation of fluorescence of extract from n=3 yeast transformants.

The current model for Pex1/Pex6 function in yeast peroxisome biology posits that the assembled ATPase heterohexamer is recruited to the peroxisomal membrane by Pex6 binding to the tail-anchored protein Pex15 [Birschmann et al 2003]. Once recruited to the peroxisomal membrane, Pex1/Pex6 are thought to increase the import efficiency of proteins into the peroxisomal lumen by recycling Pex5, the receptor for the C-terminal peroxisomal targeting signal (PTS1) [Terlecky et al 1995, Dodt et al 1995]. In the cytosol, Pex5 binds the PTS1 tag on client proteins and subsequently docks at the peroxisome membrane through interactions with Pex13/Pex14/Pex17 [Albertini et al 1997; Huhse et al 1998; Erdmann and Blobel 1996]. The PTS1-tagged protein translocates across the membrane fully folded [Walton et al 1995; Romano et al 2019], through a dynamic pore that may consist of an oligomer of Pex13, Pex14, and Pex5 [Meinecke et al 2010; Gao et al 2022; Ravindran et al 2023]. During translocation, Pex5 enters the peroxisome lumen, ultimately presenting only the N-terminus to the cytosol [Gouviea et al 2003; Skowyra et al 2022]. A complex of membrane bound E3 ligases, Pex2/Pex10/Pex12, mono-ubiquitinates Pex5 on its N-terminus [Platta et al 2007; Platta et al 2009; Feng et al 2022]. The Pex1/Pex6 motor, recruited to the peroxisomal membrane by Pex15, is then thought to recognize and engage mono- ubiquitinated Pex5, extracting it from the peroxisome membrane [Platta et al 2005]. After de- ubiquitination by Ubp15 [Debelyy et al 2011], the cytosolic, de-ubiquitinated Pex5 can repeat the cycle for additional rounds of matrix-protein import.

This model of Pex1/Pex6 activity is reminiscent of that of Cdc48 in ER-associated degradation, where Cdc48 extracts terminally misfolded proteins from the endoplasmic reticulum after ubiquitination by membrane bound E3 ligases [Schliebs, Girzalsky, Erdmann 2010]. Both Cdc48 and Pex1/Pex6 use a threading mechanism for substrate processing, in which ‘pore loops’ with conserved aromatic residues in the central ATPase channel make steric contact with the substrate polypeptide, and hydrolysis-driven vertical movements of the ATPase subunits pull the substrate through the central pore of the hexamer [Bodnar et al 2017, Gardner et al 2018]. Notably, only the D2 ATPase ring in Pex1/Pex6 actively hydrolyzes ATP, while both the D1 and D2 ATPase rings in Cdc48 are active ATPases. Both Cdc48 and Pex1/Pex6 are recruited to the target membrane by anchored adaptor proteins, Ubx2 for Cdc48 and Pex15 for Pex1/Pex6 [Schuberth and Buchberger 2005, Neuber et al 2005]. Both motors are thought to recognize their substrates through a ubiquitin modification, though Cdc48 engages a large repertoire of poly-ubiquitinated substrates, while Pex1/Pex6 may engage a relatively limited set of mono-ubiquitinated substrates (Pex5 and Pex5-like receptors, Pex18 and Pex21). It remains unclear whether Pex1/Pex6 can engage other substrates, such as Atg36 [Yu et al 2022], and how Pex1/Pex6 selectively recognizes mono-ubiquitinated Pex5, presumably through its poorly characterized N-terminal domains.

Pex1 and Pex6 each have two N-terminal domains (N1 and N2, **Figure 1A**) that are homologous to Cdc48’s single N-terminal domain [Blok et al 2015]. Only the N1 domain of Pex1 is well conserved at the sequence level. Despite the poor conservation, structure predictions suggested that Pex1 N2 and both N domains of Pex6 are related to the N-terminal domain of Cdc48 and NSF (**Figure 1A**). Cryo and negative stain electron microscopy (EM) [Blok et al 2015, Gardner et al 2015, Ciniawsky et al 2015] show that the Pex6 N1 domain sits equatorial to the D1 ATPase ring, and the Pex1 N2 and Pex6 N2 domains are located above the D1 AAA-ATPase ring. While Pex1 N1 has been previously crystallized in isolation [Shiozawa et al 2004], it is not observed in EM structures, presumably because it is flexibly tethered to Pex1 N2 and does not occupy a specific location relative to the ATPase rings. Unlike the N-terminal domains of NSF and Cdc48, none of the Pex1 or Pex6 N-terminal domains have been observed to undergo conformational changes that correlate with nucleotide hydrolysis in the ATPase domains [Gardner et al 2015, Ciniawsky et al 2015].

Previous work in yeast demonstrated that Pex15 binding to Pex1/Pex6 is essential for peroxisome function. Without Pex1, Pex6, or Pex15, ubiquitinated Pex5 accumulates at the peroxisome membrane [Platta et al 2005; Kragt et al 2005] and peroxisomes are targeted for autophagy [Nuttall et al 2014; Yu et al 2022]. Pex15 is typically regarded as a ‘tether’ required to recruit Pex1/Pex6 to the peroxisomal membrane. However, there are also indications for additional roles, for instance, co-immunoprecipitation experiments suggested that Pex15 bridges the Pex1/Pex6 and Pex5/Pex14 complexes [Gardner et al 2018, Tamura et al 2014]. *In vitro*, the cytosolic domain of Pex15 (Pex15 1-309) is a substrate of Pex1/Pex6 and inhibits its ATPase activity in a pore-loop dependent manner, but it is unclear whether the ATPase inhibition or Pex15 unfolding is relevant *in vivo*, where Pex15’s C-terminus is tethered to the membrane (Gardner et al 2018]. Negative-stain EM of the Pex1/Pex6/Pex15 complex showed that the core α-helical region of Pex15 binds near the Pex6 N2 domain [Gardner et al 2018], which is in agreement with *in vivo* interaction studies showing that Pex15 binds optimally to the Pex6 N-terminal domains and D1 ring [Birschmann et al 2003]. However, the low resolution of this structure obscured any details of the interaction between Pex15 and Pex6, and the role of the Pex6 N1 domain in Pex15 interactions remained unclear. Intrigued by the functional importance of the Pex15/Pex6 interaction, despite its poor sequence conservation, and the unique equatorial position of the Pex6 N1 domain, we set out to investigate the role of the Pex6 N1 domain in Pex1/Pex6 motor function.

Here we show that the Pex6 N1 domain is essential for Pex1/Pex6 function at the peroxisome in *S. cerevisiae*. An X-ray crystal structure of the isolated N1 domain from yeast Pex6 showed that despite limited conservation of primary sequence, this domain shares the conserved structure of the NSF and Cdc48 N terminal domains. When docking this structure of the isolated Pex6 N1 domain into a cryo-EM reconstruction of the Pex1/Pex6 complex, we found that there is additional density unaccounted for by the atomic models. Based on models generated with AlphaFold-Multimer, we hypothesized that this additional density arises from a stable contact between Pex6’s N1 domain and an extension of the Pex1 D2 ATPase domain, which we confirmed through crosslinking assays. This contact helps stabilize the correct alternating assembly of the Pex1/Pex6 heterohexamer. We also found that the Pex6 N1 domain is required for binding to Pex15 *in vitro*. Given its ability to interact with Pex15 and the active D2 ATPase domain, the Pex6 N1 domain is uniquely positioned to coordinate co-factor binding and ATP-hydrolysis activity.

## Results

To test whether the Pex6-N1 domain is essential for Pex1/Pex6 function in *S. cerevisiae*, we first assessed the active hexamer formation of Pex1/ΔN1-Pex6 *in vitro*. We isolated recombinant Pex1/ΔN1-Pex6, in which Pex6 lacks the first 184 amino acids, by a two-step affinity purification via a His tag on the N-terminus of Pex6 and a C-terminal FLAG tag on Pex1 **(Figure 1B)**. Using an enzyme-coupled ATPase assay, we found that the Pex1/ΔN1-Pex6 motor had ATPase activity comparable to wildtype Pex1/Pex6, indicating that the Pex1/Pex6 AAA-ATPase can form an active ATPase in the absence of the Pex6 N1 domain **(Figure 1C)**.

To test if the Pex6 N1 domain is essential for Pex1/Pex6’s role in peroxisomal matrix protein import *in vivo*, we used a colorimetric assay based on the incorporation of a synthetic pathway for converting tryptophan into the fluorescent pigment prodeoxyviolacein (PDV) [Deloache et al 2016]. The last enzyme in the pathway, VioE, is tagged with the C-terminal peroxisome-targeting signal, such that the enzyme is sequestered to peroxisomes in wildtype cells, but remains in the cytosol of cells defective for peroxisome import. Thus, yeast defective in peroxisome import produce PDV, which is detectable by its green pigmentation and fluorescence. Transformation of a Δ*pex6* strain with a plasmid expressing wildtype Pex6 recovered peroxisome import, while a plasmid expressing ΔN1-Pex6 did not, indicating that the N1 domain is essential for Pex1/Pex6 function in matrix protein import **(Figure 1D)**.

To further characterize the Pex6 N1 domain, we recombinantly expressed the Pex6 1-184 fragment, which was chosen based on its stability in limited proteolysis assays **(Supplementary** Figure 1A). Pex6 1-184 was crystallized, and we collected X-ray diffraction data to a resolution of 1.9 Angstroms **(Supplementary** Figure 1B, **Table 1)**. We then used molecular replacement with an AlphaFold atomic model to determine the structure of the Pex6 N1 domain **(Figure 2A)**. The Pex6 N1-domain consists of two subdomains: a double-psi *β* barrel in the N-terminal half (N1a) followed by an *α/β* roll subdomain (N1b). Despite low sequence identity, the domain is structurally similar to the N-domain of the related AAA-ATPase Cdc48 **(Figure 2B)**.

**Figure 2.**
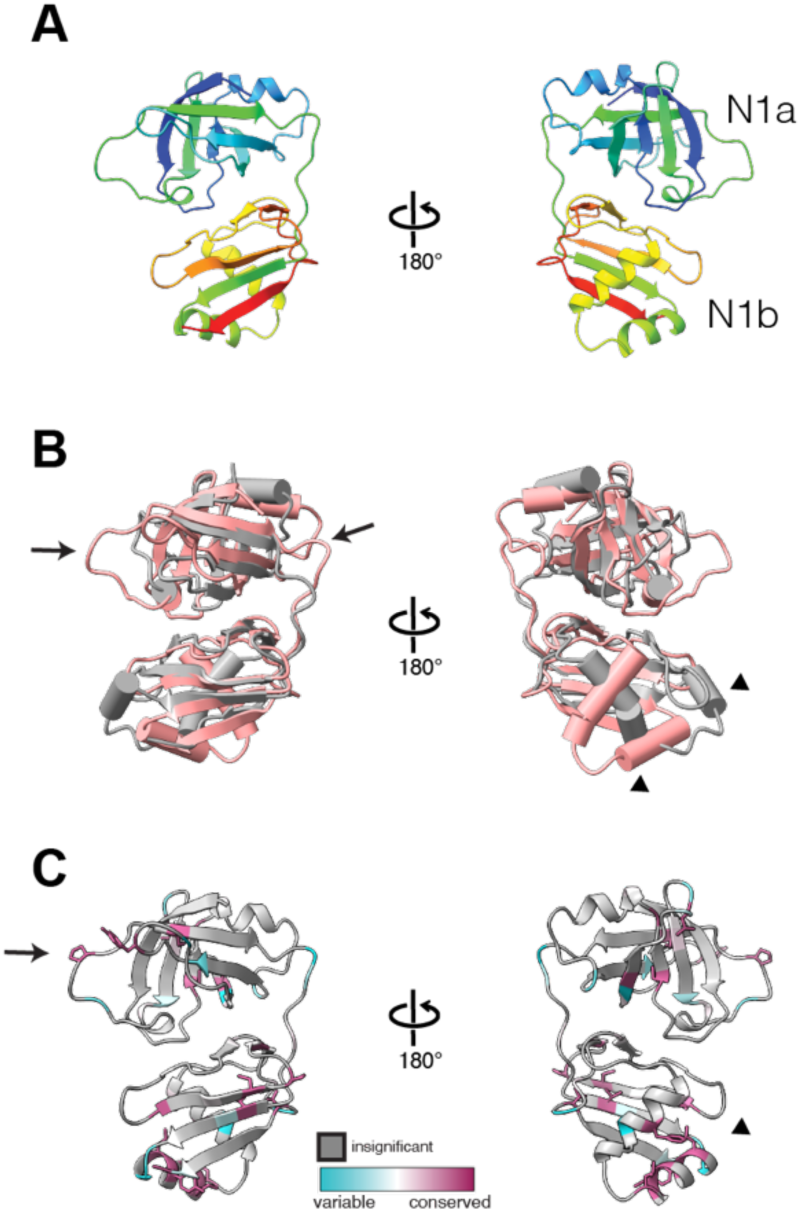
A) Atomic model of yeast Pex6 N1 domain resolved by X-ray crystallography, cartoon colored from N (blue) to C terminus (red). B) Overlay of Cdc48 N domain (gray, PDB 4KDL) and Pex6 N1 domain (pink). C) ConSurf analysis of conserved residues in the Pex6 N1 domain.

**Table 1.**
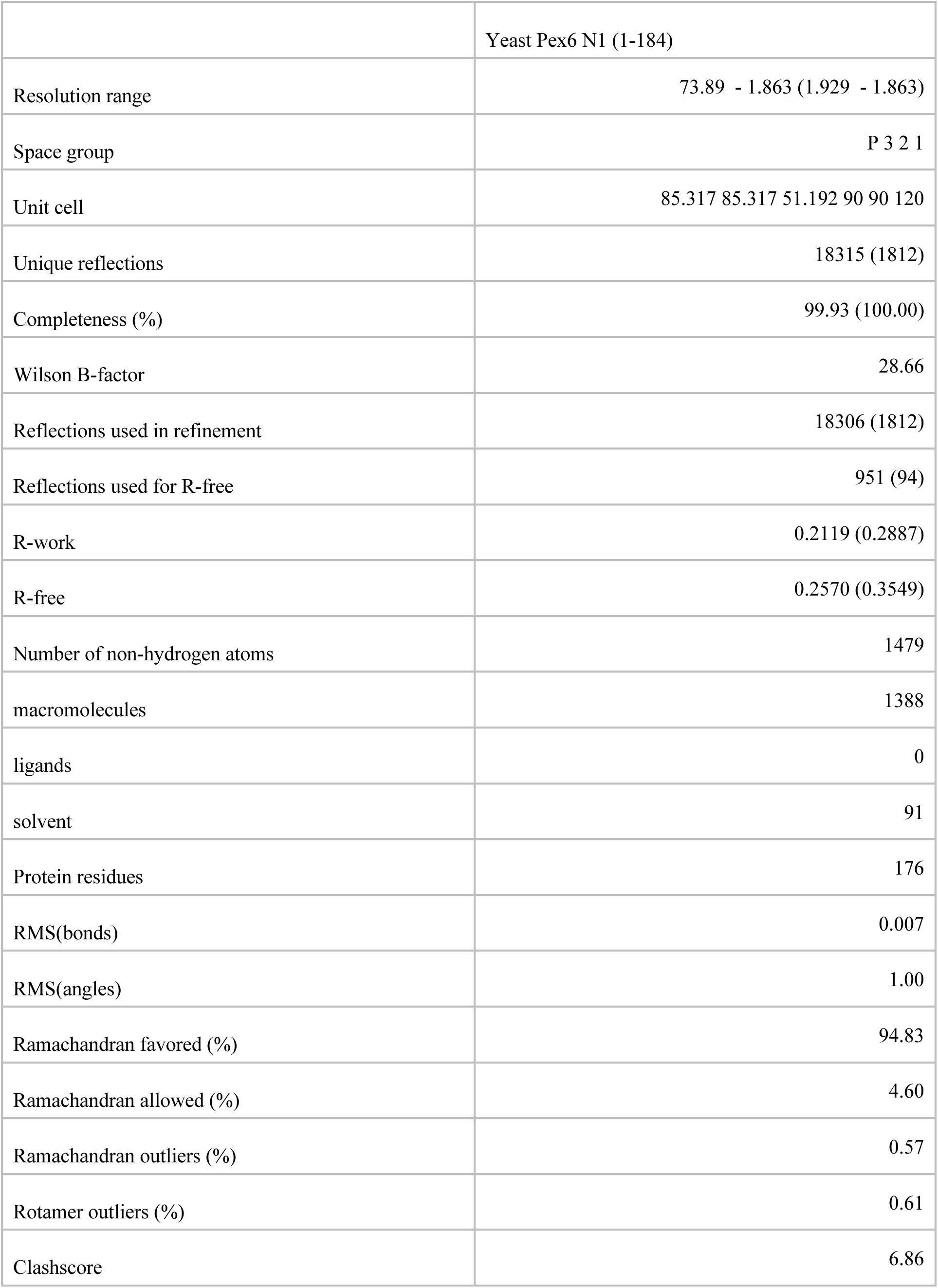
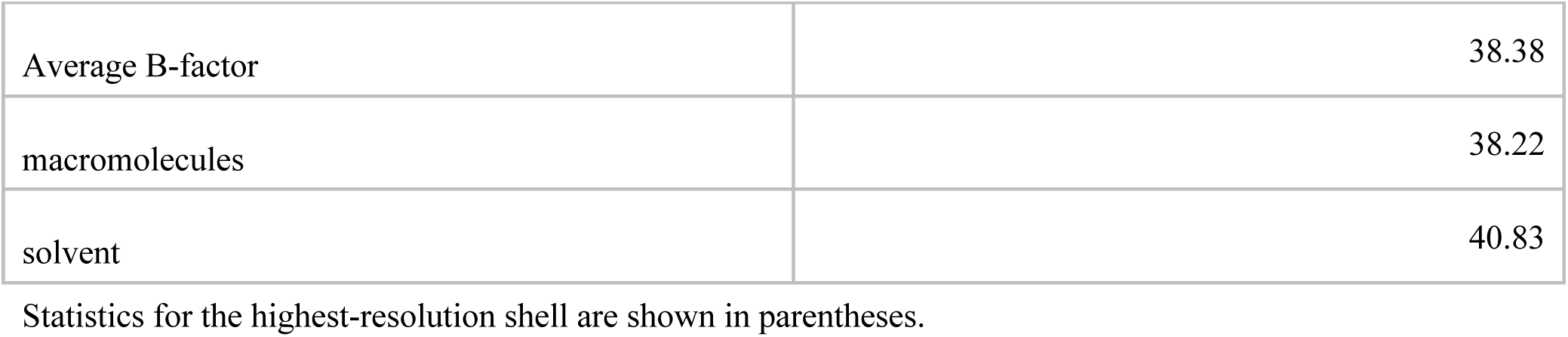
Data Collection and Refinement Statistics.

|  | Yeast Pex6 N1 (1-184) |
| --- | --- |
| Resolution range | 73.89 - 1.863 (1.929 - 1.863) |
| Space group | P 3 2 1 |
| Unit cell | 85.317 85.317 51.192 90 90 120 |
| Unique reflections | 18315 (1812) |
| Completeness (%) | 99.93 (100.00) |
| Wilson B-factor | 28.66 |
| Reflections used in refinement | 18306 (1812) |
| Reflections used for R-free | 951 (94) |
| R-work | 0.2119 (0.2887) |
| R-free | 0.2570 (0.3549) |
| Number of non-hydrogen atoms | 1479 |
| macromolecules | 1388 |
| ligands | 0 |
| solvent | 91 |
| Protein residues | 176 |
| RMS(bonds) | 0.007 |
| RMS(angles) | 1.00 |
| Ramachandran favored (%) | 94.83 |
| Ramachandran allowed (%) | 4.60 |
| Ramachandran outliers (%) | 0.57 |
| Rotamer outliers (%) | 0.61 |
| Clashscore | 6.86 |
| Average B-factor | 38.38 |
| macromolecules | 38.22 |
| solvent | 40.83 |

Statistics for the highest-resolution shell are shown in parentheses.

Structure-based alignment of the Pex6 N1 domain and Cdc48 N domain shows that Pex6 N1 has more prominent loops in the N1a subdomain **(arrows, Figure 2B)** and an altered position of the N1b helices **(arrowheads, Figure 2B)**. The Pex6 N1 domain surface is largely hydrophilic (**Supplementary** Figure 2A) and has a negatively charged surface in the N1a subdomain and positively charged patch in the N1b subdomain (**Supplementary** Figure 2B). Using the seven homologous sequences identified by ConSurf for this isolated domain, we mapped the location of the most conserved residues on the Pex6 N1 domain structure (**Figure 2C**). In addition to more highly conserved residues in the hydrophobic core of the N1a and N1b subdomains, we observed conserved residues in the extended loops of the N1a subdomain (**Figure 2C, arrow**) and a cluster of residues between the N1b helices that form a small hydrophobic patch (**Figure 2C, arrowhead**).

To understand the role of the Pex6 N1 domain in the Pex1/Pex6 hexamer, we analyzed the Pex6 N1 domain structure in the context of a structure of recombinant yeast Pex1/Pex6 determined by cryo-electron microscopy. This Pex1/Pex6 reconstruction was determined in the presence of ATP and an *in vitro* substrate mEOS-Pex15 1-309 (**Figure 3A, B**) and allowed building models for Pex1/Pex6’s D1 and D2 ATPase rings at ∼4 angstrom and ∼5-9 angstrom resolution, respectively (**Supplementary** Figure 3). The overall architecture is consistent with Pex1/Pex6 structures previously determined by negative stain and cryo-electron microscopy [Blok et al 2015, Gardner et al 2015, Ciniawsky et al 2015]. We docked and rebuilt AlphaFold models of Pex1 and Pex6 into the cryo-EM map, starting with the Pex1 and Pex6 subunits at the highest overall resolution (Pex6 – chain B and Pex1 – chain C), and then fitting these models into the density of the other subunits. Although our biochemical assays suggest that Pex1/Pex6 unfolds the Pex15 1-309 domain of the mEOS-Pex15 1-309 fusion before stalling on the mEOS moiety (**Supplementary** Figure 4 **and Supporting Information**), we did not observe density attributable to substrate in the central pore of this structure, which was readily apparent from the open central pore in both 2D class averages and the final 3D reconstruction. Note that we did not expect to observe mEOS-Pex15 1-309 bound to Pex6 N-terminal domains, akin to the Pex1/Pex6/Pex15 complex in EMD-7005, as Pex15 1-309 is engaged by the ATPase as a substrate, unfolded, and likely dislodged from the N domains by motor-mediated translocation.

**Figure 3.**
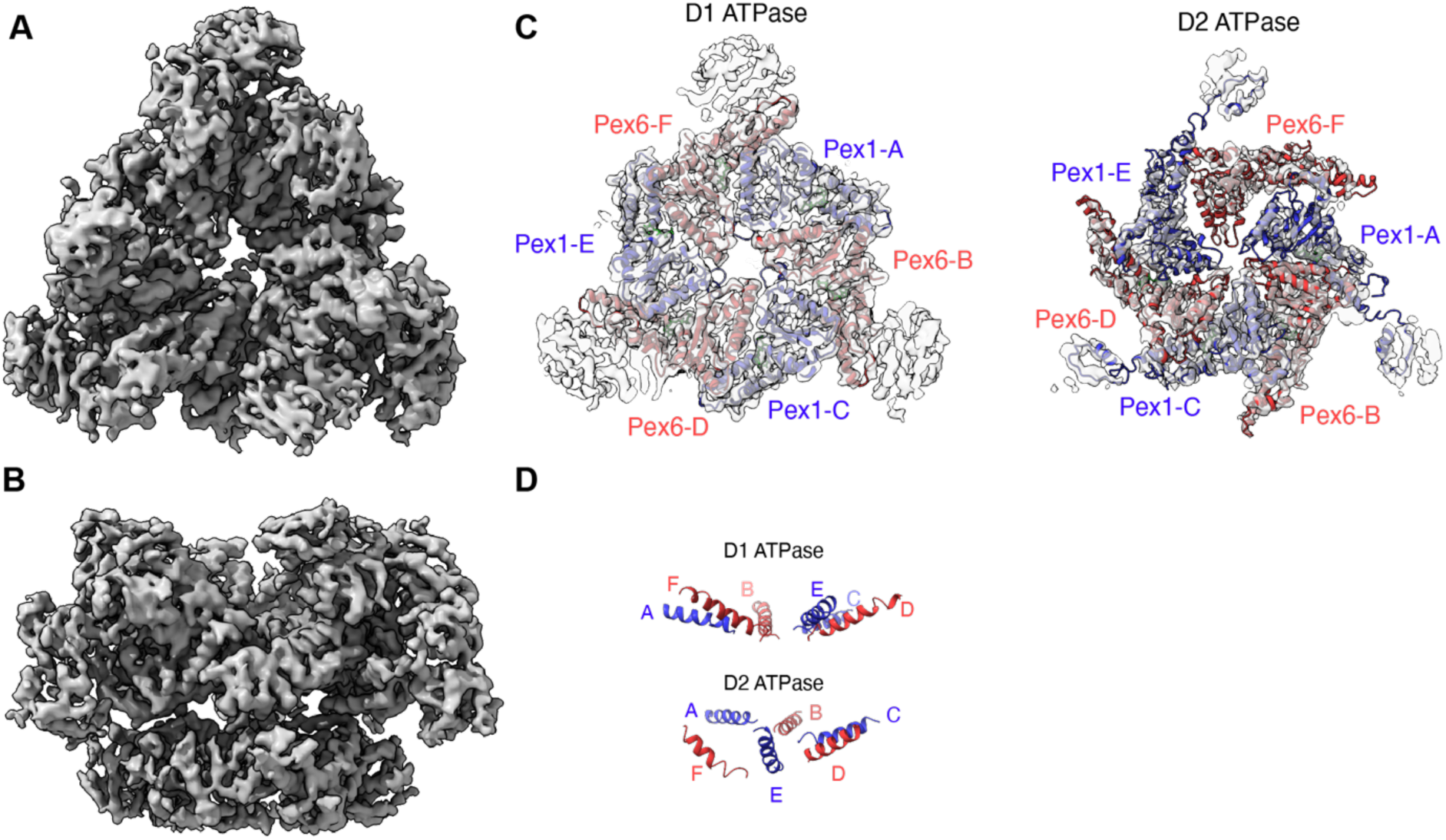
A) Top view of cryo-EM structure of Pex1/Pex6 in ATP. B) Sideview of cryo-EM structure of Pex1/Pex6 in ATP. C) Top views of slices showing the D1 ATPase ring and D2 ATPase ring, with atomic models for Pex1 (blue) and Pex6 (red). D) Side view of the relative positions of Pex1 and Pex6 pore loop α-helices for the D1 ATPase domains (Pex1 499-517, Pex6 518-539) and D2 ATPase domains (Pex1 772-789; Pex6 807-822) showing a planar arrangement for the D1 ring and a “spiral staircase” for the D2 ring.

In a prototypical Type 2 AAA-ATPase, the nucleotide binding sites are located at the interfaces between subunits and contribute to both motor assembly and activity. The ATPase domain of one subunit binds the nucleotide, contributing both the characteristic Walker A motif, required for nucleotide binding, and the Walker B motif, required for nucleotide hydrolysis. The adjacent subunit contributes to the same site by interacting with the nucleotide through its arginine finger motif, which is required for nucleotide hydrolysis. Conformational changes arising from nucleotide binding and hydrolysis coordinate the ATP-hydrolysis steps in adjacent subunits and in the motor complex. Given the heterohexameric architecture of the Pex1/Pex6 complex, there are 4 unique nucleotide binding sites: both the Pex1 and Pex6 nucleotide binding sites in the D1 ring, and the Pex1 and Pex6 nucleotide binding sites in the D2 ring. Both the Pex1 and Pex6 D1 domains harbor endogenous substitutions in the motifs required for ATP hydrolysis, while the D2 domains are well conserved. Therefore, the D1 ATPase ring has been hypothesized to bind, but not hydrolyze ATP, and to serve as a stable scaffold for motor assembly, while nucleotide hydrolysis in the D2 ATPase ring is essential for substrate processing *in vitro* and Pex1/Pex6 function *in vivo*.

For the cryo-EM reconstruction, Pex1/Pex6 and mEOS-Pex15 1-309 were incubated in the presence of ATP, which should lead to active motor states with ATP- and ADP-bound subunits. Consistent with the hypothesis that the D1 ATPase ring acts as a stable scaffold for motor assembly, we find density for nucleotide in all the D1 ATPase domains, which we modeled as ATP. Both the Pex1 and Pex6 D1 ATPase interfaces are stabilized by contacts between the small AAA subdomain and the large AAA subdomain of the adjacent protomer, particularly anchored by the hydrophobic residues Pex1 Y654 and Pex1 I451, as well as additional points of contact between large AAA subdomains near the central pore (**Supplementary** Figure 5). While the D1 ATPase domains do not have conserved hydrophobic pore loops, the helices that would position these loops in the central pore are on the same plane, making the D1 ATPase ring planar and symmetrical (**Figure 3D**). In contrast to the well resolved D1 ATPase ring, the resolution of the D2 ATPase domains is lower and varies by subunit position. This is consistent with a vertical “spiral staircase” configuration that has been previously observed for other AAA-ATPases [Gates and Martin 2020]. In this spiral staircase, subunit Pex1-A is in the highest position, Pex1-E is in the lowest position, and Pex6-F is the “seam” subunit at an intermediate position (**Figure 3D**). The density and positions of Pex1-A, Pex6-B, Pex1-C, and Pex6-D are consistent with these subunits being nucleotide bound, and we have therefore modeled them with ATP in the binding pocket. Both Pex1-E and Pex6-F are at a lower resolution than the other domains and farther from the adjacent subunit and therefore were not modeled with nucleotide bound. The position of the D1 ATPase domains does not appear to be coordinated with the nucleotide occupancy or vertical position of the corresponding ATPase domain in the spiral staircase of the D2 ring, consistent with the D1 ATPase ring maintaining the assembly of Pex1/Pex6.

The atomic models derived from the crystal structure of the isolated Pex6 N1 domain aligned very well with the atomic model based on the cryo-EM map (RMSD 1.604) (**Figure 4A**). The biggest differences in the models were related to the loops with high B-factors in the crystal structure, which form the interface of the N1 domain with the N2 domain of Pex6 (**Figure 4B**). For example, two loops in the N1a subdomain (aa 40-50 and aa 74-83) form contacts with the Pex6 N2 domain utilizing conserved residues Tyr43 and Pro80 (**Figure 4B**).

**Figure 4.**
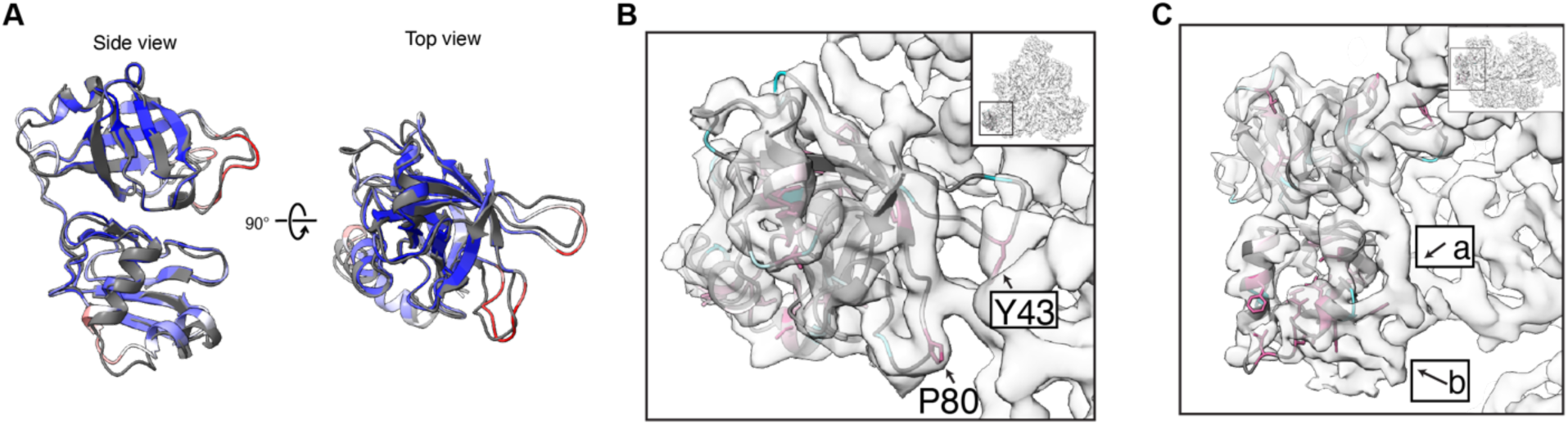
A) Alignment of the Pex6 N1 atomic models derived from the X-ray crystal structure of the isolated Pex6 N1 (colored by B-factor) and the cryo-EM map (gray), RMSD = 1.604 angstroms. B) Top view of Pex6 N1with an atomic model from the cryo-EM map shown as cartoon and colored by ConSurf conservation as in Fig 2, with the same orientation as in Fig. 4A. Loops with the most variation appear to mediate the interface with the N2 domain. C) The atomic model from the X-ray crystal structure of the isolated Pex6 N1 domain (colored by ConSurf conservation) does not fill two regions of density: region a, predicted to be the Pex6 N2 domain, and region b, adjacent to conserved residues in the N1b subdomain.

Interestingly, the atomic model derived from the X-ray crystal structure of the isolated Pex6 N1 domain did not account for two regions of density in the cryo-EM map. Region ‘a’ (**Figure 4C**) is a band of density along the side of the N1 domain that appears to connect the last residue modeled in the X-ray crystal structure, residue 175, to the Pex6 N2 domain (**Figure 4C**). The AlphaFold prediction for full-length Pex6 suggests that this density corresponds to the linker between the N1 and N2 domains. Although the N1 domain construct used for crystallography included some of this linker (175-186), we could not resolve these nine C-terminal residues, likely because they are flexible in absence of the N2 domain and the remainder of Pex6.

A second region of density (‘b’, **Figure 4C**) is near the base of the N1b subdomain and adjacent to conserved residues in Pex6 N1. While previous models of Pex1/Pex6 suggested that this density corresponds to part of the linker between Pex6’s N1 and N2 domains, closer examination of ours and others’ EM structures for Pex1/Pex6 show density bridging the Pex1 D2 domain and the Pex6 N1 domain in certain motor conformations (EMD-7005, EMD-2584). We used AlphaFold-Multimer to predict the structure of Pex1 D2 in complex with Pex6’s N-terminal domains (**Figure 5A**). AlphaFold confidently predicted an interaction between Pex6 N1 domain and a loop (aa 941-967) extending from Pex1’s D2 small α-helical subdomain (**Figure 5A, Supplementary** Figure 8) that would explain the density seen in cryo-EM reconstructions of the Pex1/Pex6 hexamer. The prediction suggests that, in the assembled motor, a disordered linker extends from the Pex1 D2 ATPase to position a β-sheet and α-helix for binding to the Pex6 N1b subdomain (**Figure 5A**).

**Figure 5.**
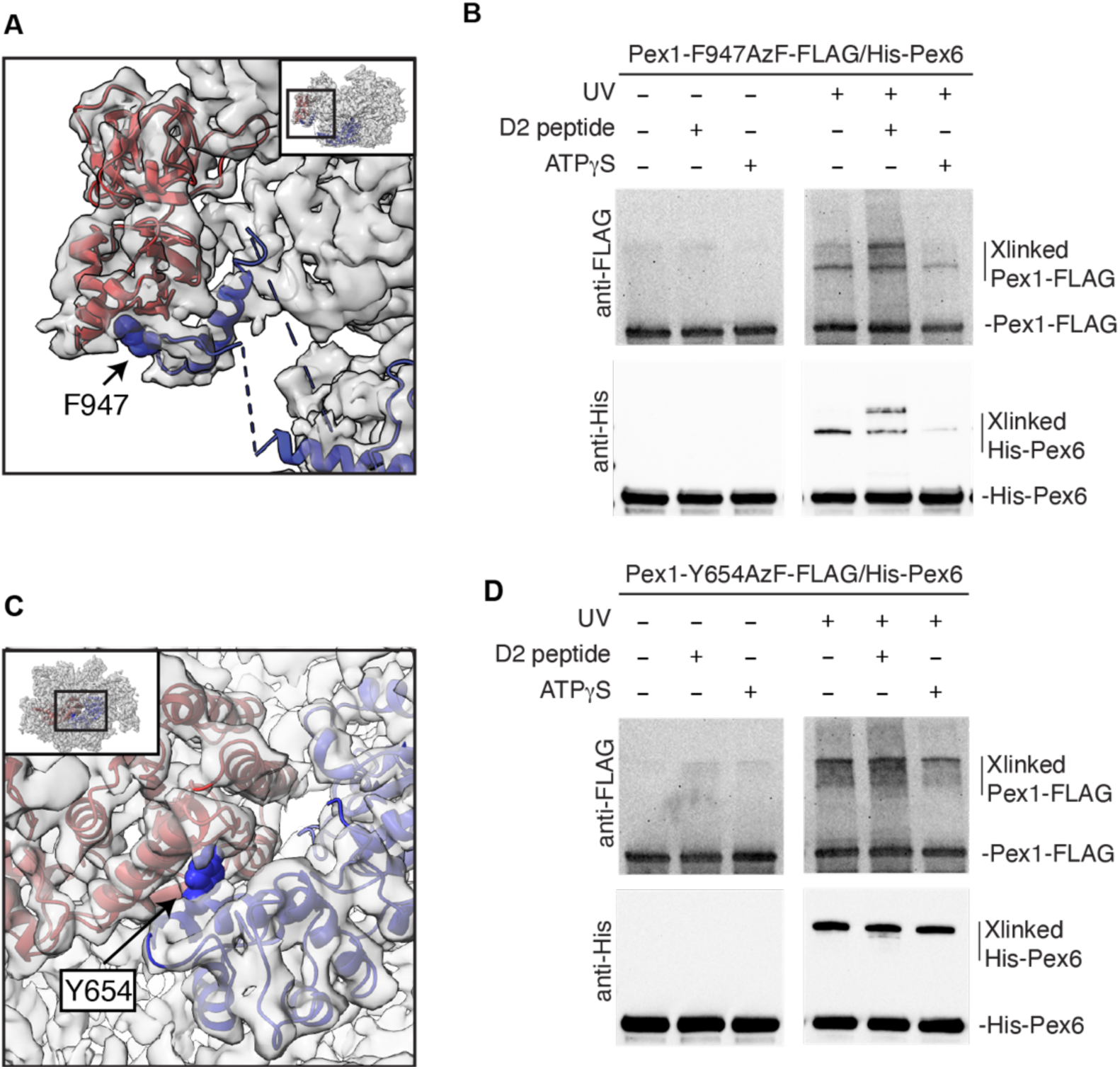
A) Modeling of the Alphafold-predicted interaction in the cryo-EM volume (gray) shows Pex1 (blue) contributing an α-helix and ý-sheet to the missing density near Pex6 N1 domain (red). F947 sidechain shown with van der Waals radii. B) UV crosslinking shows Pex1- F947AzF-FLAG crosslinks with His-tagged Pex6. The crosslinking to His-Pex6 is reduced by ATPγS and altered by a competing peptide derived from the predicted Pex1 D2 extension. C) Pex1 Y654 (sidechain shown with van der Waals radii) is at the interface between the small AAA subdomain of Pex1 D1 and the large AAA subdomain of Pex6 D1. D) Crosslinking between Pex1-Y654AzF-FLAG and His-Pex6 under the same conditions as in B.

To verify a potential interaction, we introduced a photoreactive artificial amino acid, azidophenylalanine (AzF), into Pex1 at the predicted contact site (Pex1 F947AzF) (**Figure 5A**). We recombinantly expressed Pex1-F947AzF-FLAG together with His-Pex6 and purified the complex by tandem affinity purification. As a positive control, we also purified Pex1 Y654AzF/Pex6, as Y654 mediates the interaction between the D1 small α-helical subdomain of Pex1 and the D1 large AAA subdomain of Pex6 (**Figure 5C**). Upon UV exposure, both Pex1 Y654AzF and Pex1 F947AzF crosslinked to Pex6 (**Figure 5B, D**) to produce a His and FLAG-tagged product of approximately 200 kDa, confirming that the incorporated azidophenylalanines in Pex1 contact Pex6. Addition of a chemically synthesized peptide of the Pex1 D2 loop sequence (Pex1 aa 941-967) to the crosslinking reaction causes Pex1-F947AzF to form a new Pex1/Pex6 crosslinked product, which migrates more slowly than the crosslinked product observed in the absence of this competitor peptide (**Figure 5B**). These findings are consistent with the peptide at high concentrations displacing the Pex1 D2 loop from Pex6 N1, causing it to crosslink to a new location on Pex6. We note that Pex1 Y654AzF also forms an additional crosslinked product to His-Pex6 in the presence of peptide, albeit to a lesser extent (**Figure 5D)**. This suggests that the addition of the Pex1 D2 loop peptide does not disrupt Pex1/Pex6 assembly, but may alter its conformation. These crosslinked products do not form in the presence of apyrase, which hydrolyzes ATP and ADP to AMP, indicating that the crosslinking occurs only in a nucleotide-dependent assembly of Pex1 and Pex6 **(Supplementary** Figure 6B).

Incubation with a slowly hydrolyzable nucleotide analog, ATPγS, reduced crosslinking between Pex1 F947AzF and Pex6, but had no effect on Pex1 Y654AzF, suggesting that the interaction between Pex1’s D2 loop and Pex6’s N1 domain is dependent on the nucleotide state of the motor and potentially the formation of a spiral staircase in the D2 ring with a combination of ADP-bound and ATP-bound subunits **(Figure 5B, D)**. Integrating the cryo-EM structure, AlphaFold2 prediction, and crosslinking data suggests that the extension from the Pex1 D2 small AAA subdomain contacts the Pex6 N1b subdomain, and we therefore incorporated this contact site into the atomic models for the Pex1/Pex6 complex **(Figure 6 A, B)**.

**Figure 6.**
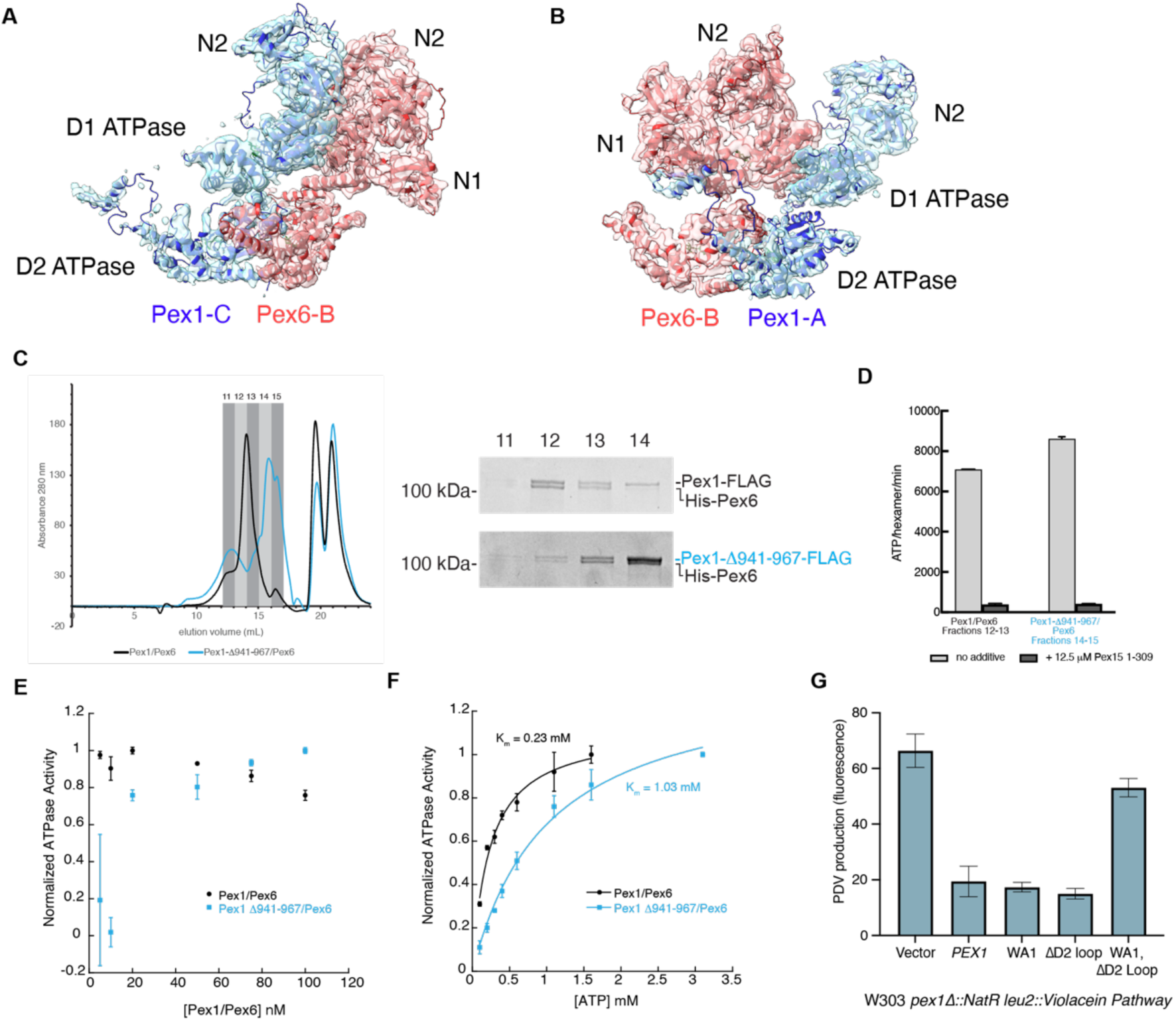
A) Atomic models of Pex1 and Pex6 at the Pex6:Pex1 ATPase interfaces refined from Alphafold models docked into the cryo-EM density. B) Same as A, but showing the Pex1:Pex6 ATPase interface. C) Pex1-ý941-967/Pex6 elutes from a Superose6i column in later fractions than Pex1/Pex6. D) Concentrated Pex1-ý941-967/Pex6 from fractions 14-15 has ATPase activity similar to WT Pex1/Pex6 concentrated from fractions 12-13. Shown are the average and standard deviation of three technical replicates. E) Titration of Pex1/Pex6 and Pex1 Δ941- 967/Pex6 motor concentration shows a decline in activity for Pex1 Δ941-967/Pex6 at low hexamer concentrations. F) Michaelis-Menten analysis of the ATP hydrolysis activity for wildtype Pex1/Pex6 and Pex1 Δ941-967/Pex6. G) The colorimetric assay for peroxisome import efficiency in yeast shows that Pex1-ý941-967 and Pex1 K467S (D1 Walker A mutation) can complement a ý*pex1* deletion strain, but Pex1-K467S- ý941-967 does not support peroxisome import.

There are two interfaces for assembly of the Pex1/Pex6 heterohexamer: the Pex6:Pex1 interface between the Pex6 ATP binding sites and neighboring Pex1 protomer (**Figure 6A**) and the Pex1:Pex6 interface that forms at the Pex1 ATP binding sites and neighboring Pex6 protomer **(Figure 6B)**. The interactions between the active D2 ATPase domains are dynamic, and thus both the Pex6:Pex1 and Pex1:Pex6 interfaces depend on nucleotide binding in the D1 ATPase domains [Gardner et al 2015] for stable heterohexamer assembly. The Pex6:Pex1 interface is also further supported by electrostatic interactions between the Pex6 and Pex1 N2 domains (**Supplementary** Figure 7A). To test if interaction between the Pex1 D2 loop and the Pex6 N1 domain contributes to Pex1/Pex6 assembly, we used a two-step affinity purification to isolate the Pex1-ý941-967/Pex6 motor. The deletion of aa 941-967 removes the extension predicted to interact with Pex6 N1. We found that Pex1-ý941-967-FLAG and His-Pex6 co-purify, yet elute in later fractions than wild-type Pex1/Pex6 from the Superose6 gel filtration column **(Figure 6C)**, suggesting that the heterohexameric assembly is destabilized. When these fractions were concentrated to encourage oligomerization and subsequently subjected to an ATPase assay, we found that the Pex1-ý941-967/Pex6 motor had near wildtype levels of ATPase activity that could be significantly inhibited by Pex15 (**Figure 6D**), consistent with re-assembly of an active heterohexamer at higher concentrations after the motor partially dissociated over the course of the purification. To confirm a contribution of the Pex6 N1:Pex1 D2 loop contact for Pex1/Pex6 hexamer stability, we measured the dependence of the ATPase activity on the concentration of the motor. For these experiments, Pex1/Pex6 was equilibrated for 5 minutes at the desired concentration of motor prior to measuring its ATPase activity. While the wildtype Pex1/Pex6 motor maintains activity even at low concentrations, the Pex1-ý941-967/Pex6 motor loses activity at low concentrations, likely due to dissociation (**Figure 6E**). Furthermore, we found that Pex1-ý941-967/Pex6 has a more than 4-fold higher K_m_ for ATP compared to wild type Pex1/Pex6, requiring higher concentrations of ATP to reach maximal activity (**Figure 6F)**.

To test if the deletion of the Pex1 D2 loop affected Pex1/Pex6 motor function *in vivo*, we used the colorimetric assay for peroxisome import and determined the ability of the Pex1-Δ941- 967 to complement Δ*pex1* in yeast. We found that Pex1-Δ941-967 complemented the *pex1* deletion similar to wildtype Pex1, suggesting no defect in Pex1/Pex6 assembly in these conditions. Given that the Pex1:Pex6 interface also relies on nucleotide binding in the D1 and D2 ATPase domains, we then tested Pex1-Δ941-967 in the context of a Walker A mutation in the Pex1 D1 ATPase domain. We found that Pex1-WA1 could also recover peroxisome import in PDV-synthesizing yeast similar to wild type Pex1 (**Figure 6G**), indicating that neither nucleotide binding in the Pex1 D1 domain nor the D2 loop:Pex6 N1 interaction are required for Pex1/Pex6 function *in vivo* under our test conditions. However, a Pex1 mutant with both the WA1 mutation and the Δ941-967 deletion exhibited a defect in peroxisomal import, indicating that the nucleotide binding in the D1 and the Pex6 N1:Pex1 D2 loop interaction additively support Pex1:Pex6 assembly. Together with the *in vitro* data that Pex1-ý941-967/Pex6 motor has a higher K_m_ for ATP than wildtype Pex1/Pex6, these results suggest that interactions between the D1 domains of Pex1 and Pex6 as well as the contacts between Pex6 N1 and Pex1 D2 loop interaction both critically contribute to heterohexamer stability, with the Pex6 N1:Pex1 D2 loop interaction becoming more important at low ATP concentrations.

We observed that deletion of the Pex6 N1 domain hinders Pex1/Pex6 function at the peroxisome *in vivo* (**Figure 1C**). Given that the extension of the Pex1 D2 ATPase domain is not required for efficient peroxisomal import under our growth conditions, Pex6 N1’s contribution to Pex1/Pex6 heterohexamer stability is unlikely to explain the peroxisomal import defect of the N1 domain deletion. Since we had previously observed an interaction between the purified Pex6 N1 domain and the cytosolic domain of Pex15 in immunoprecipitation experiments [Gardner et al 2018), we next tested if the N1 domain is essential for Pex1/Pex6 binding to Pex15 by measuring the response of Pex1/ΔN1-Pex6’s ATPase activity to the presence of the Pex15 1-309 cytosolic domain. Unlike WT Pex1/Pex6, we found that the ATPase activity of Pex1/ΔN1-Pex6 was not inhibited by Pex15 1-309, indicating that the Pex6 N1 domain is essential for Pex15 binding and/or engagement by the ATPase motor **(Figure 7A)**.

**Figure 7.**
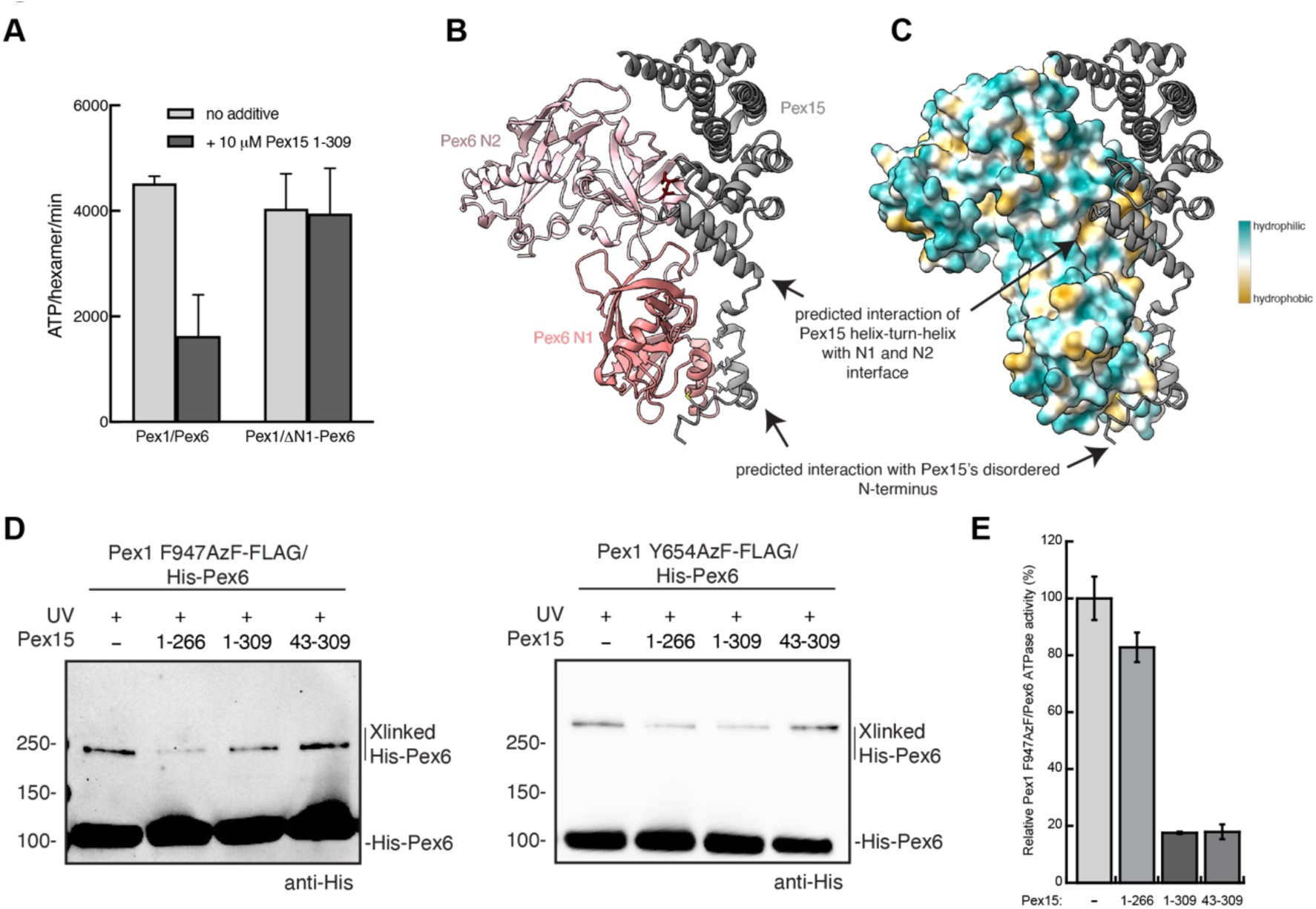
A) The ATPase activity of Pex1/ΔN1-Pex6 is not inhibited by the cytosolic domain of Pex15 (Pex15 1-309). Shown are the average and standard deviation for three technical replicates. B) Alphafold-Multimer prediction of interactions between the Pex15 core and N-terminus (gray) and the Pex6 N1 (salmon) and N2 (light pink) domains. Pex6 N1 is predicted to bind Pex15’s disordered N-terminus and also the first helix of the core domain. Residues in red were previously shown to be protected from solvent by Pex15 binding (Gardner et al 2018). C) Alphafold prediction of interactions between the Pex15 core and N-terminus (gray) and the Pex6 N1 and N2 domains (surface representation colored by lipophilicity). D) Efficiency of Pex1 F947AzF and Pex1 Y654AzF crosslinking to His Pex6 in the absence or presence of Pex15 1- 309, Pex15 1-266 (lacking C-terminal tail for pore loop engagement), and Pex15 43-309 (lacking N-terminal region for binding to the N1b subdomain). E) Effect of Pex15 constructs on Pex1 F947AzF/Pex6 ATPase activity normalized to the ATPase activity in the absence of Pex15. Shown are the averages and standard deviations for three technical replicates.

To model the potential interaction between Pex15 and the Pex6 N-terminal domains, we utilized AlphaFold-Multimer **(Figure 7B, C)**. In this prediction, the core domain of Pex15 arches over Pex6’s N2 domain, with a helix-turn-helix motif of Pex15 (aa 43-69) forming contacts in the cleft between the N1 and N2 domains. This prediction is consistent with our previous finding in hydrogen-deuterium exchange experiments that a peptide in this N-domain cleft is protected from solvent by Pex15 binding (**Figure 7B**, red, Gardner et al 2018]. AlphaFold-Multimer also predicts that the disordered N-terminus of Pex15 interacts with the Pex6 N1b subdomain (**Figure 7 B,C**). In this prediction, leucine residues near the Pex15 N terminus bind at a conserved hydrophobic patch on the N1b subdomain **(Figure 2C)**. In support of this interaction, we previously found that N-terminal truncations or mutations that remove these leucine residues in the disordered N-terminal region of Pex15 reduce the apparent affinity for Pex1/Pex6 [Gardner et al 2018]. We conclude that the Pex6 N1 domain is required for Pex15 binding to Pex1/Pex6, making contact with Pex15’s disordered N terminus and interacting with Pex15’s globular domain at the N1-N2-domain interface. Therefore, the loss of binding to Pex15 is likely the reason why ΔN1-Pex6 does not complement a ý*pex6* deletion in yeast.

According to these predictions, both the disordered N terminus of Pex15 and the Pex1 D2 loop interact with the Pex6 N1b subdomain. To determine whether Pex15 binding influences the conformation of the Pex1 D2 loop, we analyzed the crosslinking efficiency between Pex1 F947AzF and His-tagged Pex6 in the presence of Pex15’s cytosolic domain. We found that both Pex15 1-309 and Pex15 1-266 reduced the crosslinking efficiency between Pex1 F947AzF and His-tagged Pex6 (**Figure 7D**). The inhibition of Pex1 F947AzF/Pex6 crosslinking depended on the disordered N-terminus of Pex15 that is predicted to bind the Pex6 N1b subdomain, as Pex15 43-309 did not show the same inhibitory effect as as Pex15 1-309 (**Figure 7D**). However, this crosslinking inhibition did not correlate with the level of ATPase inhibition, as Pex15 43-309 and Pex15 1-309 reduced ATP hydrolysis to a similar extent in these conditions, whereas Pex15 1-266 showed only minimal ATPase inhibition (**Figure 7E**). Crosslinking between Pex1 Y654AzF/Pex6 was also slightly reduced by Pex15 (**Figure 7D**), indicating that Pex15 binding influences the conformation of the Pex1:Pex6 interface in general. Pex15 binding to the Pex6 N-terminal domains may thus alter the interaction with the Pex1 D2 loop and enable an allosteric mechanism that links Pex1/Pex6 docking with Pex15 at the peroxisome membrane to ATP hydrolysis in the D2 ATPase ring.

## Discussion

Here we demonstrated that the Pex6 N1 domain contributes to Pex1/Pex6 function by binding the peroxisomal membrane receptor Pex15 and by stabilizing the Pex1:Pex6 interface through the interactions with a loop extending from the Pex1 D2 ATPase domain. We found that the Pex6 N1 domain is essential for motor function *in vivo*, while deletion of the Pex1 D2 loop is only detrimental in the context of additional hexamer destabilization through a Pex1 D1 Walker A mutation. Given that deletion of the Pex6 N1 domain dramatically reduces the affinity of Pex1/Pex6 for Pex15 *in vitro*, and reciprocal Pex15 mutations that reduce Pex1/Pex6 affinity were shown to cause peroxisomal import defects, [Gardner et al 2018], we propose an essential role for the Pex6 N1 domain in binding Pex15 and thereby recruiting Pex1/Pex6 to the peroxisomal membrane. Additionally, the contribution of the Pex6 N1:Pex1 D2 loop interaction to the hexamer stability may be essential under certain growth conditions, for instance, at low Pex1/Pex6 concentrations or reduced cellular levels of ATP.

The dual role of Pex6 N1 in binding to the peroxisomal tether Pex15 and the Pex1 D2 ATPase domain raises the question of whether these binding events influence one another and play a role in tuning Pex1/Pex6 function at the peroxisome. Intriguingly, we found that the addition of Pex15’s cytosolic domain reduced crosslinking between Pex1 F947AzF and His- Pex6. This suggests that Pex15 binding, and thus peroxisomal membrane tethering, alters the conformation of the Pex1:Pex6 interface through an allosteric mechanism mediated by Pex6 N1 domain. Negative stain EM of the Pex1/Pex6/Pex15 complex in the presence of ATP shows Pex15 bound to all three Pex6 subunits and density for a connection between the Pex6 N1 domain and Pex1 D2 ATPase domain for two of the three Pex1 ATPases (EMDB-7005) [Gardner et al 2018]. These data suggest that Pex15 binding to Pex6 N1 does not preclude the contact with the Pex1 D2 loop, and may stabilize an interaction beyond the Pex1 941-967 region. Based on these data, we hypothesize that there are several different possible conformations for the interaction between the Pex1 D2 and Pex6 N1 that may be modulated by Pex15 binding. Such conformational changes could allow for preferential stabilization of the heterohexamer, presentation of the Pex1 D2 loop for recognition by another protein, or alteration of Pex1 D2 ATPase dynamics when bound by Pex15 at the peroxisomal membrane.

Despite the poor conservation of the Pex6 N1 domain and Pex15, our findings have intriguing implications for the role of the PEX6 N1 domain in metazoa. Pex15 has functional orthologs, PEX26 and APEM9, in metazoa and plants. Similar to Pex15, human PEX26 is predicted to be a tail-anchored α-helical protein that was shown to interact with the PEX6 N-terminal domains [Tamura et al 2014; Furuki et al 2006]. Indeed, AlphaFold-Multimer also predicts an interaction between PEX26 and Pex6 N-terminal domains in humans and Arabidopsis [Judy et al 2022]. Intriguingly, AlphaFold2 predicts that the PEX6 N1 domain in humans is partially disordered in the absence of PEX26, but folds into the conserved structure in the presence of PEX26 [Judy et al 2022]. Thus, not only is the interaction between the PEX6 N1 and PEX26 likely conserved in metazoa, it may play an important role in stabilizing the fold of the PEX6 N1 domain.

There is also evidence for a function of the D2 loop in human PEX1/PEX6. This extension of the small AAA subdomain of PEX1’s D2 is much longer in the human (∼150 residues) versus the yeast homolog (∼50 residues). The length and predicted α-helix in the midst of the otherwise disordered loop are well conserved, and we therefore hypothesize that similar to the Pex1 D2 loop in yeast, this extended PEX1 D2 loop in humans also influences motor assembly or regulation through interactions with the PEX6 N domains. The dissociation of PEX1/PEX6 is commonly observed in Peroxisome Biogenesis Disorders arising from mutations in PEX1 or PEX6, thus mechanisms that improve PEX1/PEX6 assembly are of potential therapeutic interest.

## Experimental Procedures

### Yeast strains

All yeast strains are derived from a parental background of W303 MATa ura3-1 his3-11 trp1-1 leu2-3 leu2-112 can1-100. Individual knockout strains (pex1Δ::NatR, pex6Δ::NatR) were constructed using standard transformation techniques with pFA6A-NatMX, previously described in Gardner et al 2018.

### Tandem Affinity Purification of Pex1-FLAG/His-Pex6

Recombinant yeast Pex1-FLAG/His-Pex6 complexes were purified as previously described in Gardner et al 2018. Briefly, BL21*(DE3) *E. coli* co-expressing pCOLADuet Pex1- FLAG and pETDuet His-Pex6 were grown from overnight cultures until OD600 = 0.6. Pex1 and Pex6 expression were induced with 0.3 mM IPTG for an overnight induction at 18°C. Bacteria were harvested at 4000 RPM for 15 minutes and resuspended in NiA buffer (50 mM HEPES pH 7.6, 100 mM NaCl, 100 mM KCl, 10% glycerol, 10 mM MgCl2, 0.5 mM EDTA, 20 mM imidazole, 1 mM ATP) with protease inhibitors (PMSF, aprotinin, leupeptin, pepstatin) and 1 mg/mL lysozyme. Resuspended cells were lysed by sonication, then centrifuged at 30,000xg for 30 minutes, and the supernatant was batch bound to Ni-NTA resin. Bound proteins were eluted in buffer with an additional 500 mM imidazole supplemented with 1 mM ATP and the elution was applied to an anti-FLAG agarose column. FLAG-tagged proteins were eluted in GF buffer (60 mM HEPES pH 7.4, 100 mM NaCl, 100 mM KCl, 5% glycerol, 10 mM MgCl2, 0.5 mM EDTA, 1 mM ATP) with 0.15 mg/mL FLAG peptide and concentrated with a 100 MWCO spin concentrator, then snap-frozen in liquid nitrogen and stored at -80°C. Hexameric Pex1/Pex6 complexes were further purified by size exclusion chromatography using a Superose6 Increase 10/300 column equilibrated in GF buffer with 1 mM ATP.

### Enzyme-coupled ATPase assay

The ATPase activity of Pex1-FLAG/His-Pex6 was monitored using an ATP/NADH coupled enzyme assay that uses the production of NADH as a functional readout for hydrolysis of ATP. Typical reactions contain 3 U mL^−1^ pyruvate kinase, 3 U mL^−1^ lactate dehydrogenase, 1 mM NADH, 7.5 mM phosphoenolpyruvate, 5 mM ATP, and 2 uM BSA with variable concentrations of Pex1/Pex6 heterohexamer (typically 5 nM-20 nM) and Pex15 substrates (typically 1 uM - 5 uM). The absorbance of NADH was measured at 340 nm in a 96-well plate on a SpectraMax plate reader for 15 minutes at 30 °C.

### Colorimetric Assays for Peroxisomal Import

The pex1Δ::NatR and pex6Δ::NatR knockout strains with the incorporated violacein pathway were made by integrating pWCD1401 or pWCD1402 at the leu2-3 locus as previously described [Gardner et al 2018]. Complementation by wildtype or mutant Pex1 or Pex6 was determined by transformation with CEN ARS plasmids with the PEX1pr-PEX1 variant or PEX6pr-PEX6 variant. Mutations were incorporated by Gibson cloning or around the horn PCR of the vector backbone and blunt-end ligation of the phosphorylated ends.

The CEN ARS plasmids with PEX1 and PEX6 variants were transformed into the knockout, violacein- pathway containing yeast strains and selected on SD-Ura-Leu. Three colonies from each transformation were streaked for singles on SD-Ura-Leu plates, then a single colony from each was picked for growth overnight in 5 mL YPD media at 30° and 180 RPM. For visualization of the green pigmentation, yeast were spotted on SD-Ura-Leu plates (5 uL, OD600=0.3) and grown at 30 degrees C. For quantification of pigmentation by fluorescence, yeast were then diluted 1:50 into 5 mL of selective SD-Ura-Leu media and grown for 60 hours at 30°C and 180 RPM. 800 uL of culture were spun down and resuspended in 150 uL of glacial acetic acid and transferred to thin-walled PCR tubes. The samples were heated to 95°C for 15 minutes, mixed by inverting, then incubated for an additional 15 minutes at 95°C. The product was transferred to Eppendorf tubes and centrifuged for 5 minutes at 4700 RPM, then 100 uL of supernatant was vacuum filtered through 0.2 um PTFE filters in a 96-well plate format (Pall corporation). Fluorescent intensity at an excitation wavelength of 535 nm and an emission wavelength of 585 nm was measured.

### Purification of Pex6 N1 1-184

His-PP-Pex6 N1 constructs were recombinantly expressed in BL21* E coli grown in DYT from plasmid pDC73 or pDC69. Cultures were grown shaking at 37°C and were induced at ∼OD_600_ = 0.6 by the addition of IPTG (C_f_ 0.3 mM), then incubated shaking for 4.5 hrs at 30 °C. Cells were harvested by centrifugation at 6,000 x *g* for 20 minutes at 4°C, then resuspended in Ni_A buffer (25 mM HEPES pH 7.6, 100 mM NaCl, 100 mM KCl, 10% glycerol, 10 mM MgCl_2_, 0.5 mM EDTA, 20 mM imidazole) supplemented with protease inhibitors: leupeptin, pepstatin, aprotinin, 2 mg/mL lysozyme, and benzonase (Novagen). Resuspended cell pellets were stored at -80 °C until thawed for purification. Thawed cells were sonicated on ice for 2 min, and the lysate was clarified by centrifugation for 30 minutes at 30,000 x *g*. The subsequent soluble extract was batch bound to Ni-NTA resin for 30 minutes, rotating end-over-end. The protein-bound resin was washed with Ni_A buffer for ∼40 CVs. His-PP-Pex6 N1 was eluted with 10 CVs of Ni_B buffer (25 mM HEPES pH 7.6, 100 mM NaCl, 100 mM KCl, 10 % glycerol, 500 mM imidazole). The affinity purification tags were cleaved by PreScission protease overnight while dialyzing into Ni_A buffer. Uncleaved protein His-PP-Pex6 N1 and PreScission Protease were removed by incubation with Ni-NTA resin. The resulting flow-through was concentrated and run on a HiLoad 16/60 Superdex 200 column (GE Life Sciences) into crystallization buffer: 20 mM HEPES pH 7.6, 50 mM NaCl, 50 mM KCl, 0.5 mM TCEP.

### Limited Proteolysis

His-PP-Pex6 1-204 protein (purified by recombinant expression from plasmid pDC69) was digested with 0.1 or 0.2 μg/μL trypsin protease (Sigma) for 10 minutes at 23 °C in 100 mM Tris pH 8.0 buffer. An aliquot of the reaction was quenched with the addition of PMSF and SDS-PAGE sample buffer (12.5 mM Tris pH 6.8, 10% glycerol, 2% SDS, 0.005% bromophenol blue) and run on an SDS-PAGE gel for size analysis. The remaining reaction was quenched with an equal volume of 6 M guanidinium-HCl and flash frozen until analyzed by mass spectrometry. Mass spectrometry revealed two cleavage sites at residues Pex6 K184 and K196.

### Crystallization of Pex6 N1 1-184

Crystal conditions were screened using a Mosquito liquid-handling robot (TTP Labtech) and the JCSG screens I-IV (Qiagen). Promising conditions were further screened and scaled up 10-fold to 4 μL hanging drops with a 500 μL reservoir volume. The best crystals were obtained from 4 μL hanging drops in which 2 μL of 5 mg/mL Pex6 1-184 was mixed with 2 μL of a precipitant solution containing 0.8 M LiCl, 0.1 M citric acid pH 5, 16% PEG6000. Crystals were harvested after soaking for 30 sec in a cryoprotectant solution of precipitant solution with 25% sorbitol.

Diffraction data for Pex6 N1 were collected at the ALS beamline 8.3.1 at Lawrence Berkeley National Laboratory. Data were collected at a temperature of ∼100 K using a wavelength of 1.1158 Å. The datasets were processed in the *P 3 2 1* space group using autoPROC [Vonrhein et al 2011, Kabsch 2010, Evans 2006, Evans and Murshudov 2013, Winn et al 2011] and the structure of Pex6 N1 was solved using molecular replacement with the AlphaFold2 [Jumper et al 2021] predicted structure using Phenix [Liebschner et al 2019]. The structure was further refined with a 1.9 Å resolution cutoff using Phenix and Coot [Emsley et al 2010] to an R_work_/R_free_ of 0.212/0.257. Atomic models and figures were made using ChimeraX [Pettersen et al 2021].

Analysis of conservation relied on ConSurf [Ashkenazy et al 2016; Landau et al 2005; Glaser et al 2003].

### Sample Preparation for cryo-EM

Pex1-FLAG/His-Pex6 (pBG369/372) were co-expressed in *E. coli* BL21* with induction at OD_600_ = 0.8 with C_f_ = 0.3 mM IPTG for 16 hours at 18 deg C. The cells were spun down and resuspended in Ni_A buffer with protease inhibitors, benzonase, and lysozyme. Cells were lysed by sonication, and a 30,000xg supernatant was incubated with Ni-NTA agarose. The Ni-NTA agarose was washed with Ni_A + 1 mM ATP, and bound proteins were eluted with Ni_A + 1 mM ATP + 500 mM imidazole. The elution was then incubated with anti-FLAG affinity resin, washed with Ni_A + 1 mM ATP, and bound proteins were eluted with Ni_A + 1 mM ATP + 3X FLAG peptide.

mEos3.2-Pex15 1-309-PP-FLAG-His was expressed in *E. coli* BL21* cells with induction at OD_600_ = 0.6-0.8 with C_f_ = 0.3 mM IPTG for 16 hours at 18 deg C. Cells were harvested by centrifugation at 4000 x g for 20 minutes at 10 °C, and the cell pellet was resuspended in Ni_A buffer plus protease inhibitors, lysozyme, and benzonase. Resuspended cells were lysed by sonication, and the unlysed cells and cell debris were pelleted by centrifugation at 30,000 x g for 25 minutes at 10 °C. The supernatant was incubated with Ni-NTA agarose and washed with NiA. Bound proteins were eluted from the column with NiB. To remove the FLAG and His tags, the eluate was incubated with Prescission protease during overnight dialysis into NiA buffer. Tagged and untagged proteins were separated by incubation with Ni-NTA agarose. mEOS3.2-Pex15 was concentrated in a 30 MWCO spin concentrator and aliquots snap-frozen in liquid nitrogen in aliquots before storage at -80 °C.

In preparation for cryo-EM, the Pex1-FLAG/His-Pex6 Ni-NTA eluate was mixed with pKC1 for a final concentration of 5 uM Pex1/Pex6 and 64 uM mEOS3.2-Pex15 in NiA buffer with 1 mM ATP. The Pex1/Pex6/mEOS-Pex15 complex was selected by size exclusion chromatography over a Superose6 Increase equilibrated with GF buffer (60 mM HEPES pH 7.6, 50 mM KCl, 50 mM NaCl, 5% glycerol, 10 mM MgCl_2_, 0.5 mM EDTA) and 1 mM ATP. The hexamer peak was concentrated in a 100 MWCO spin concentrator to an estimated 13 uM of hexamer and snap frozen.

### Cryo-EM Data Collection and Image Processing

Initial screening of Pex1/Pex6 complex on C-flat grids with 2μm holes and 2μm spacing (Protochips) showed limited orientation of particles in ice. A thin amorphous carbon film was floated onto the holey grids to overcome this preferred-orientation issue. After plasma-cleaning these grids in a Solarus plasma cleaner (Gatan), 5 μL of 0.1% (w/v) poly-L-lysine hydrobromide (Polysciences inc.) was applied on the carbon surface of the grid for 90 s. Excess liquid was blotted with a filter paper, followed by three successive washes with 15 μL of water at a time and then blotted to dryness. 3 μL of the 3.5 mg/mL of the Pex1/Pex6/mEOS-Pex15 complex that was incubated on ice with 1 mM ATP for 5 minutes in GF buffer, was applied on the poly-L lysine treated grids. Excess sample was manually blotted with a Whatman Grade 1 filter paper for ∼5 s, and the grid was immediately vitrified by plunge freezing in liquid-ethane slurry at –179°C. This was carried out in a custom made plunge freezing device and the entire procedure was carried out in a 4°C cold room maintained at 98% relative humidity.

Cryo-EM data were collected on a Thermo Fisher Scientific Talos Arctica transmission electron microscope operating at 200 kV. Movies were acquired using a Gatan K2 Summit direct electron detector, operated in electron counting mode applying a total dose of 54 e^−^/Å^2^ as a 48-frame dose-fractionated movie during a 12 s exposure. Data was collected at 43,000x nominal magnification (1.15 Å/pixel at the specimen level) and nominal defocus range of −0.8 to −1.8 μm.

All single-particle cryo-electron microscopy data was processed using Cryosparc 3.3 [Punjani et al 2017]. 2,064,444 particles were extracted from 3578 motion- and ctf- corrected images. One round of 2D classification was performed to reject low resolution particles and non-particles. 1,775,103 remaining particles were homogenously refined into a 30A low-pass filtered volume derived from emd_6359 to generate an initial model. Two rounds of 3D classification and one round of heterogenous refinement was performed to further sort particles between classes. The final volume was heterogenously refined from 106,005 particles and a 30A low-pass filtered volume derived from emd_6359.

### Model Building

To build an atomic model of the Pex1/Pex6 hexamer, the Alphafold predictions of *Sc*Pex1 and *Sc*Pex6 were broken into domains and rigid body fit into the sharpened 3D cryo-EM reconstruction with ChimeraX. Using this placement, the volume for each monomer was extracted with phenix.mapbox, using a 10 angstrom box cushion. Using the monomers with the best resolution, the Alphafold predictions of *Sc*Pex1 and *Sc*Pex6 were docked and rebuilt into the boxed density with phenix.dock_and_rebuild. Ramachandran outliers were corrected with Isolde and the models were subsequently refined with phenix.realspacerefine. The D2 loop was rebuilt through alignment with the Alphafold multimer prediction for the D2 loop-Pex6 N1 interaction followed by phenix.realspacerefine of both Pex1 and Pex6 monomers at the interface. The N terminal domains and D1 domains from these models for Pex1 and Pex6 were then positioned in the volume of the less resolved monomers with phenix.dock_and_rebuild, and manually linked to the D2 ATPase domain which were fit by rigid body docking. To position nucleotides, the nucleotide position was copied from an alignment of each ATPase domain with PDB 1NSF (NSF D2 ATPase ring crystallized with ATP). The combined model was manually refined in Isolde and phenix.realspacerefine. We note that the models truncate the C-terminus of Pex1, which is predicted to be unfolded.

### Azidophenylalanine incorporation and photocrosslinking

An amber stop codon was cloned into pCOLADuet-PEX1-FLAG and the plasmid was co-transformed with pETDUET His-PEX6 and a plasmid encoding tRNA and tRNA synthetase (pEvol-pAzFRS.2.t1) into BL21*(DE3) *E. coli*. pEvol-pAzFRS.2.t1 was a gift from Farren Isaacs [Amiram et al 2017] (Addgene plasmid # 73546 ; http://n2t.net/addgene:73546 ; RRID:Addgene_73546). Pex1 and Pex6 were purified as described above, with the additional step that at OD600 = 0.3 tRNA synthetase expression was induced with 0.02% arabinose and azidophenylalanine (Chem-Impex) was added to C_f_ = 1 mM.

Photocrosslinking was performed in a clear, round-bottom 96-well plate with 10 uL reaction volumes. Typical reactions included 100 nM Pex1/Pex6, 10 μM BSA, 5 mM ATP or ATPγS or ATP regeneration mix, 125 μM Pex1:941-967 peptide (Elim Biopharmaceuticals Inc, FAM-NIEYFSINEHGRREENRLRLKTLLQQD) or 10 μM Pex15. The mixture was incubated on ice for 10 minutes, then exposed to UV light (Spectrolinker XL-1000, 365 nm, approximately 1500 μW/cm^2^) for 15 minutes or kept in the dark. The product was combined with 2x sample buffer (125 mM Tris pH 6.8, 20% glycerol, 4% SDS, 0.01% bromophenol blue, 10% 2- mercaptoethanol) and boiled for 5 minutes at 95°C. The samples were then used for gel electrophoresis, transferred to PVDF membranes and blotted with anti-FLAG (Sigma, F3165, 1:1000) or anti-His (Cell Signaling #2366 His-Tag 27E8) primary antibodies, then with HRP-conjugated anti-mouse secondary antibody (BioRAD 1706516) and visualized with chemiluminescence. For the apyrase controls, crosslinking of Pex1-AzF was performed after a 10 min 30 deg C incubation in ATP regeneration mix with and without apyrase (NEB#M0398S, 500units/mL, 62.5units/mL final concentration).

## AlphaFold2-multimer predictions

AlphaFold2-multimer jobs were run with ColabFold [Mirdita et al 2022; Evans et al 2021] with PDB70 templates and 3 of recycling. Statistics for the models are shown in **Supplementary** Figure 8.

## Data Availability

The cryo-EM 3D reconstruction and atomic model has been deposited with EMDB (EMD- 41788) and PDB 8U0V. The X-ray crystallography has been deposited with the PDB (PDB 8U0X).

## Acknowledgements

We thank Michal Olszewski for Cdc48 purification constructs and Ken Dong for assistance with crystallizing Pex6 N1 domain. We thank James Holton for assistance collecting and processing X-ray crystallography data. Beamline 8.3.1 at the Advanced Light Source is operated by the University of California at San Francisco with generous grants from the National Institutes of Health (R01 GM124149 for technology development and P30 GM124169 for beamline operations), and the Integrated Diffraction Analysis Technologies program of the US Department of Energy Office of Biological and Environmental Research. The Advanced Light Source (Berkeley, CA) is a national user facility operated by Lawrence Berkeley National Laboratory on behalf of the US Department of Energy under contract number DE-AC02-05CH11231, Office of Basic Energy Sciences. Bashir Ali is supported by an NSF Graduate Research Fellowship. Ryan Judy was supported by summer undergraduate research fellowship from UC Santa Barbara’s College of Creative Studies. Brooke Gardner acknowledges support from K99/R00GM121880, R35GM146784, and the Searle Scholars Program. Gabe Lander acknowledges support from R01NS095892. Andreas Martin acknowledges support of the Howard Hughes Medical Institute and R01GM094497. The content is solely the responsibility of the authors and does not necessarily represent the official views of the National Institutes of Health.

## Conflict of Interest

The authors declare that they have no conflicts of interest with the contents of this article.

## Notes

### Competing Interest Statement

The authors have declared no competing interest.

