## Supplementary Methods and Figures for "The Pex6 N1 domain is required for Pex15 binding and proper assembly with Pex1"

**Supporting Information:**

**Discussion of mEOS-Pex15 1-309 substrate**

We previously showed using hydrogen-deuterium exchange coupled with mass spectrometry that Pex1/Pex6 can unfold both the maltose binding protein domain and Pex15 domain of a MBP-Pex15 1-309 chimera [Gardner et al 2018]. To determine if substrate processing could be monitored by fluorescence, we fused the fluorescent protein mEOS3.2 to the N-terminus of Pex15 1-309. Upon exposure to light at 405 nm, mEOS3.2 undergoes backbone cleavage converting from a green fluorescent protein to a cleaved, red fluorescent protein that cannot spontaneously re-fold [Zhang et al 2021; Olszewski et al 2019]. Both mEOS3.2-Pex15 1-309 and mEOS3.2-Pex15 1-309-ssrA, which has a C-terminal tail ssrA allowing for engagement with $Cdc48^{YY/\Delta N}$ [Rothballer et al 2007], inhibited Pex1/Pex6 ATPase activity to greater extent than Pex15 1-309 (**Supplementary Figure 4A**). When we monitored unfolding by detecting the accessibility of cysteine residues to fluorescein maleimide, we found the Pex1/Pex6 unfolded the mEOS3.2-Pex15 1-309 construct. We note that there are 10 cysteines in mEOS3.2-Pex15 1-309, with seven in Pex15 1-309 and three in mEOS3.2, and we have previously monitored unfolding of Pex15 1-309 using this method (**Supplementary Figure 4B**). When we monitored unfolding by the loss of fluorescence of photocleaved mEOS3.2-Pex15 1-309-ssrA, we found that while $Cdc48^{YY/\Delta N}$ could unfold red mEOS3.2-Pex15 1-309-ssrA, Pex1/Pex6 could not (**Supplementary Figure 4C**). From these results we concluded that Pex1/Pex6 unfolds the α-helical Pex15 1-309 domain, exposing cysteine residues, but lacks the pulling force to unfold the mEOS3.2 β-barrel for loss of fluorescence.

**Supplementary Experimental Procedures**

**Maleimide labeling-based unfolding assay of Pex15**

Motor-mediated unfolding of Pex15 was monitored by the accessibility of buried cysteine residues to fluorescein-5-maleimide as previously described [Gardner et al, 2018]. mEos3.2-Pex15 1-309 and mEos3.2-Pex15-ssrA substrates were used at a final concentration of 1 μM by diluting in Buffer 1 (50 mM HEPES, 50 mM NaCl, 50 mM KCl, 10 mM $MgCl_2$, pH 7.5). The substrate was continuously maleimide-labeled by addition of F5M at a final concentration of 1 mM, and the subsequent addition of WT or mutant Pex1/Pex6 at 1 μM to the reaction mixture. The reaction was incubated for 2 minutes after the addition of Pex1/Pex6 before quenching with an equal volume (25 μL) of SDS-PAGE buffer (2% SDS, 10% β-mercaptoethanol) for a total of 50 μL reaction volume after quenching. The maximal labeling of solvent-exposed cysteines was estimated by diluting substrate in Buffer 3 (50 mM HEPES, 50 mM NaCl, 50 mM KCl, 10 mM MgCl2, 8 M urea, pH 7.5). Basal labeling of solvent-exposed cysteines was determined by incubating 1 μM substrate and 1 mM F5M in Buffer 1, without any other additives. Samples were resolved by gel electrophoresis using 10% acrylamide SDS-PAGE gel exposed to a current of 140 V for 70 minutes. Gels were imaged using Blue Epi illumination (ex: 488 nm, em: 530 nm ± 28 nm) on a Bio Rad ChemiDoc and then stained with Coomassie Blue.

**Motor-mediated unfolding of fluorescent protein mEos3.2**

mEos-Pex15 constructs were photocleaved by excitation using a 405 nm laser. The sample was added to 3.0 mm glass cuvettes on ice. Beam power was set to 0.0 mW, controlled by the pycoscope program, to avoid unnecessary usage while positioning the cuvette window towards the beam path. Power was then set to 50 mW for 10 x 2 minute intervals with sample mixing and incubation on ice between intervals. Protein was transferred to a 1.6 mL microfuge tube and snap-frozen in liquid nitrogen before storage at -80 ˚C. Laser-based excitation gave ~45% cleavage.

Unfolding of photocleaved, red mEos3.2-Pex15 1-309 and mEos-Pex15 1-309-ssrA was monitored by the change of fluorescence signal (ex: 572 nm, em: 582 nm) on a PTI spectrofluorometer using 3 nm slits (1 nm = 0.25 bandpass) controlled by FelixGX software. 2 μM photocleaved substrate and 6X ATP regeneration mix (1X: 5 mM ATP, 0.032 mg/mL creatine kinase, 16 mM creatine phosphate) in GF buffer were pre-incubated in a water bath at 30 ˚C for 10 minutes before addition to a 1.5 mm glass cuvette and monitored for 5 minutes before addition of either 0.5 uM Pex1/Pex6 or 0.5 uM $Cdc48^{YY/\Delta N}$. Motor proteins (Pex1/Pex6 and $Cdc48^{YY/\Delta N}$) were incubated at 30 ˚C only immediately before addition to the cuvette to prevent high activity rates and consumption of ATP. The substrate fluorescence was monitored for an additional 15 minutes after the addition of motor.

Supplementary Figure 1. A) SDS-P analsysis of limited proteolysis of His-Pex6 N1 1-215 revealed a stable proteolytic fragment. Mass spectrometry showed two fragments corresponding to trypsin cleavage sites at K184 and K196. B) Crystals of Pex6 1-184 grown in 1 M LiCl, 0.1 M citric acid, pH 5.0, 20% PEG6000.

**Supplementary Figure 1.**

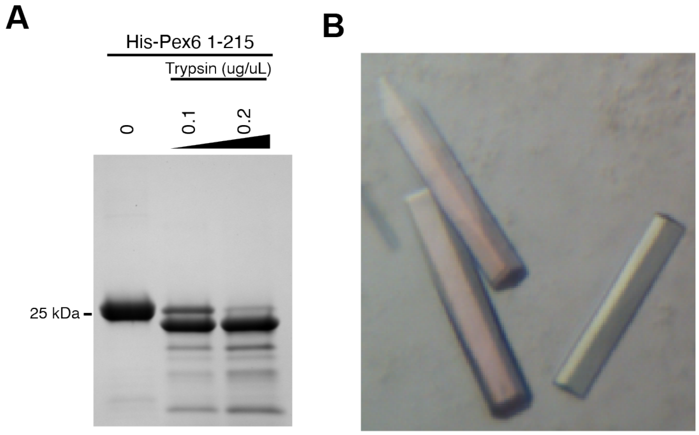

Supplementary Figure 2. A) Lipophilicity of Pex6 N1 domain, positioned as in Figure 2A. Teal – hydrophilic, yellow – hydrophobic. B) Electrostatic potential of Pex6 N1 domain, positioned as in Figure 2A. Red – acidic, blue – basic.

**Supplementary Figure 2.**

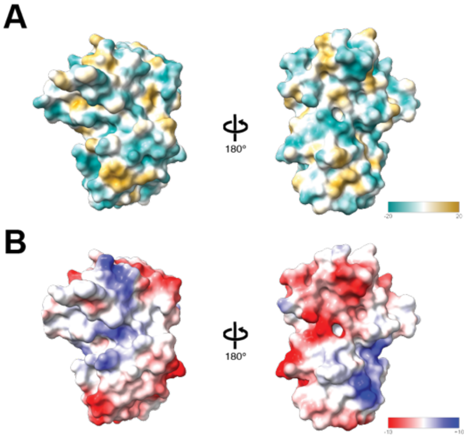

Supplementary Figure 3. A) Top, side, and bottom views of the cryo-EM reconstruction for Pex1/Pex6 in ATP with volume colored by local resolution.

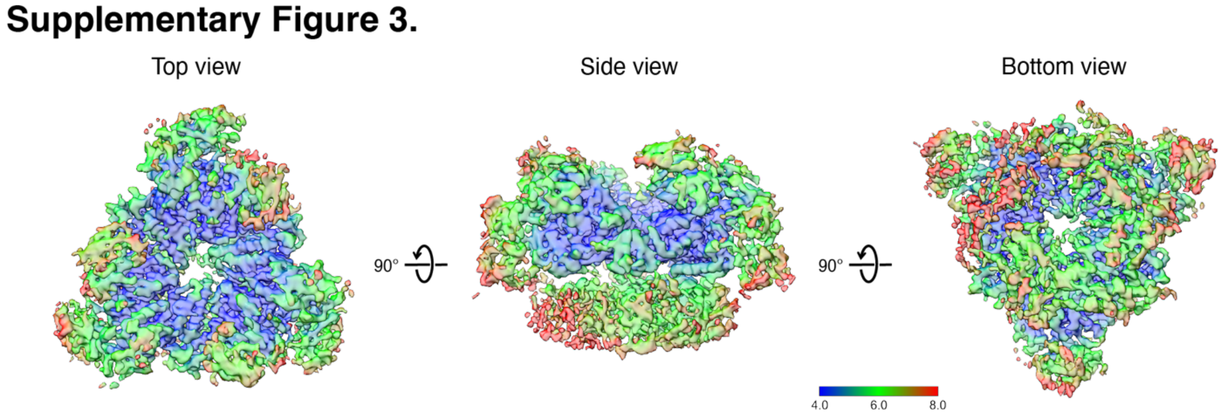

Supplementary Figure 4. A) Pex1/Pex6 ATPase activity is inhibited by Pex15 1-309, mEos-Pex15 1-309, and mEOS-Pex15 1-309-ssrA. ssrA is a C-terminal sequence of 11 unstructured residues that allows for engagement with related ATPase Cdc48. Average and standard deviation of three replicates. B) Pex1/Pex6 processing of mEos-Pex15 1-309 exposes cysteines to fluorescein-maleimide labeling, indicating pore-loop dependent unfolding. PL = Pex1-F771A/Pex6-Y805A pore loop mutant. C) Pex1/Pex6 does not alter the fluorescence of photocleaved mEOS-Pex15 1-309 (yellow) or mEOS-Pex15 1-309-ssrA (green), indicating a falue to unfold the mEOS domain. In contrast, Cdc48 readily unfolded and reduced the fluorescence of photocleaved mEOS-Pex15-ssrA (blue). D) Fluorescence from mEOS Pex15 1-309 can be detected in Pex1/Pex6 fractions eluting from a Superose6i size exclusion column, indicating migration as a complex.

### Supplementary Figure 4.

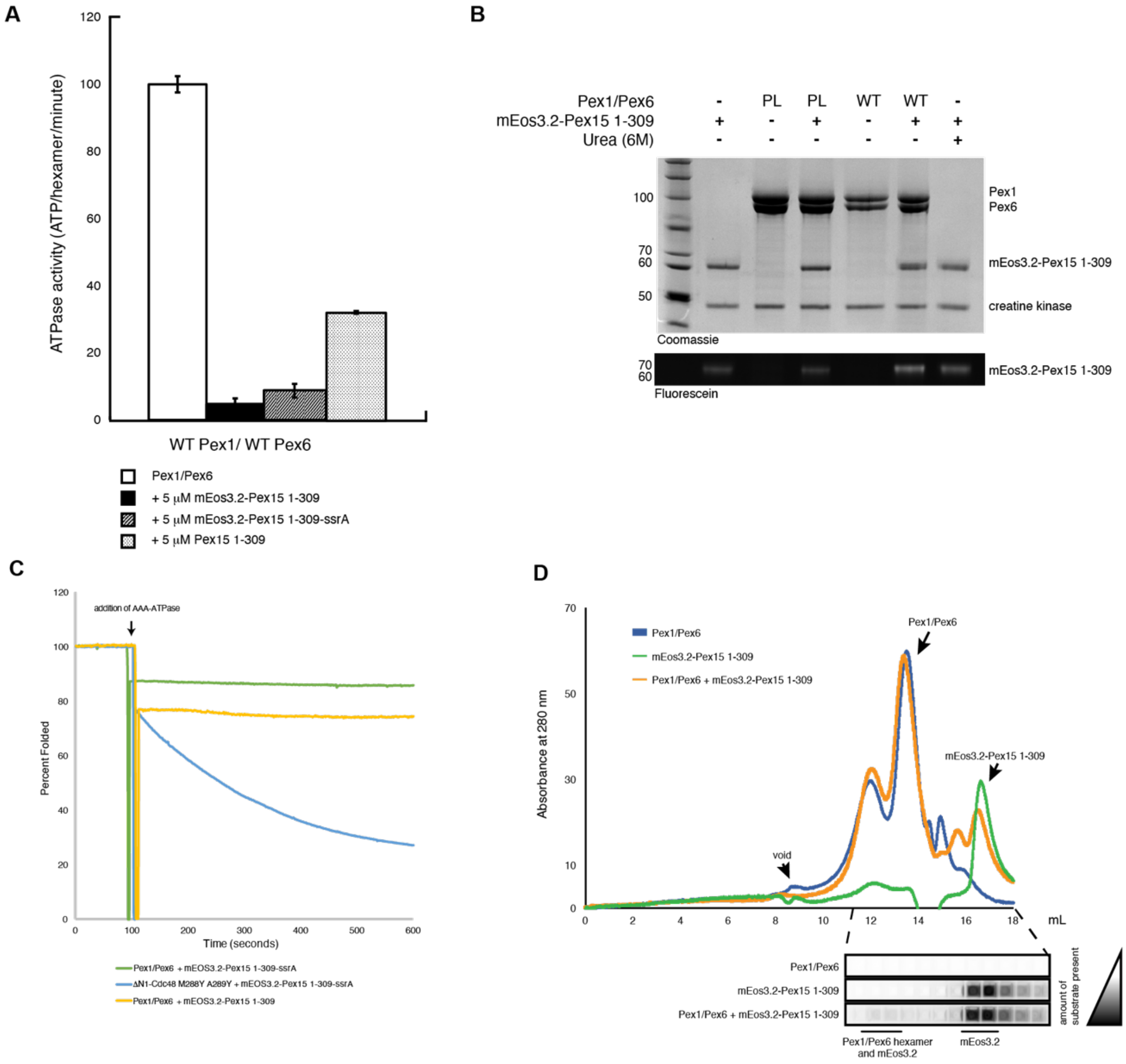

Supplementary Figure 5. A) Side view of Pex1 D1 nucleotide binding pocket. Note Pex1 Y654 at interface. B) Top view of Pex1 D1 nucleotide binding pocket.C) Side view of Pex6 D1 nucleotide binding pocket. Note Pex1 I451 at interface. D) Top view of Pex6 D1 nucleotide binding pocket.

### Supplementary Figure 5.

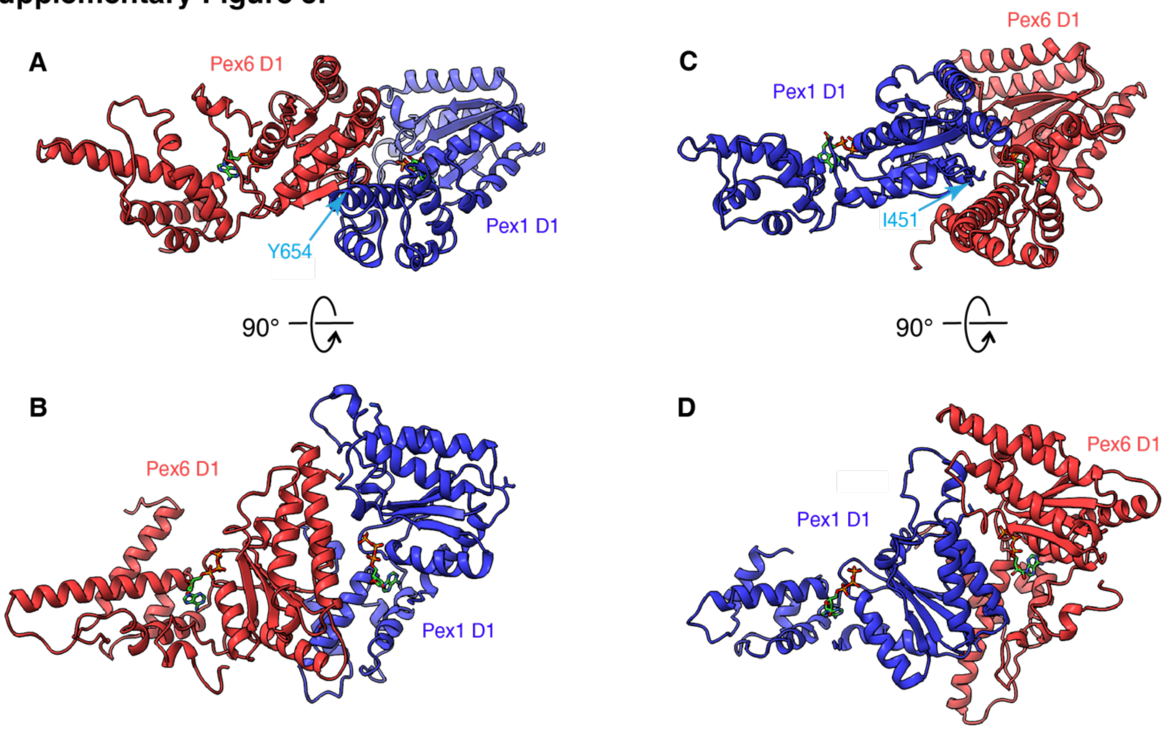

Supplementary Figure 6. A) anti-His immunoblot after SDS PAGE separation of the UV-induced crosslinking reactions between Pex1 Y654AzF-FLAG or Pex1 F947AzF-FLAG and His-Pex6 in the absence or presence of apyrase.

### Supplementary Figure 6.

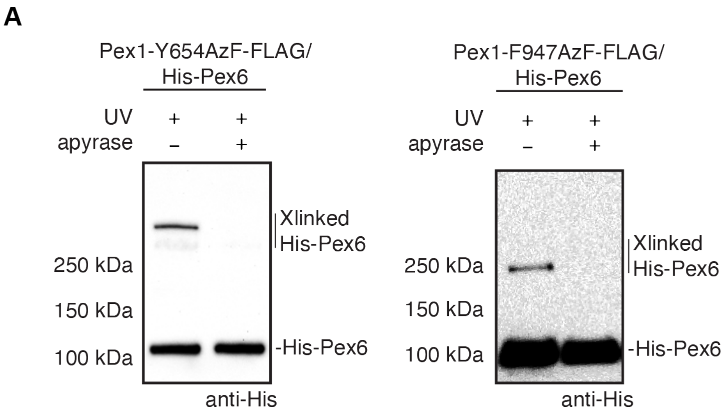

Supplementary Figure 7. A) Cartoon representation of the atomic models for Pex1 (blue) and Pex6 (red) N-terminal domains. B) Surface representation of Pex1 and Pex6 N-terminal domains in the same orientation as in A, but colored according to Coulombic potential.

### Supplementary Figure 7.

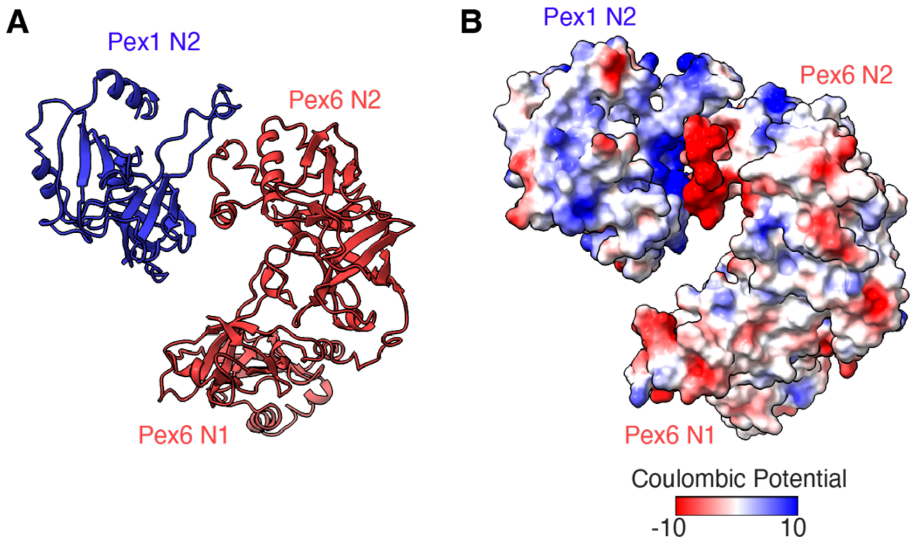

Supplementary Figure 8. Confidence metrics for AlphaFold2 predictions. Predictions were created with AlphaFold2-multimer, for which the pLLDT and pTM score is provided. For predictions with multiple unique proteins, the pLLDT-colored figures are shown twice, wherein each protein is colored by pLLDT in one structure and colored grey in the other structure.

**Supplementary Figure 8.**

| Sequences | Statistics for models in figures | Predicted aligned error | pLLDT 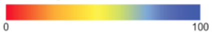  |
|---|---|---|---|
| *Sc*Pex1<br>*Sc*Pex6 N1-N2 | Figure 5.<br>mean pLLDT = 79.17<br>pTMscore = 0.56 | 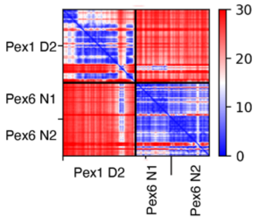  | 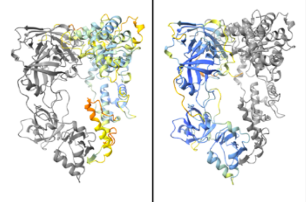  |
| *Sc*Pex15<br>*Sc*Pex6 N1-N2 | Figure 7.<br>mean pLLDT = 80.504<br>pTMscore = 0.78 | 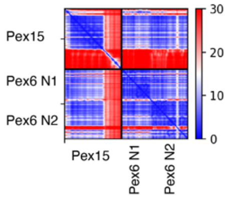  | 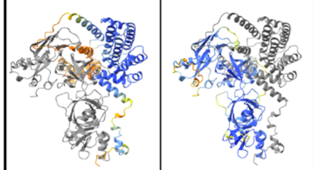  |
| *Sc*Pex1 | Figure 6.<br>mean pLLDT = 73.98<br>pTMscore = 0.59 | 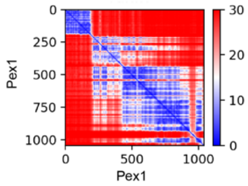  | 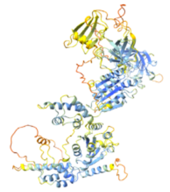 |
| *Sc*Pex6 | Figure 6.<br>mean pLLDT = 84.38<br>pTMscore = 0.80 | 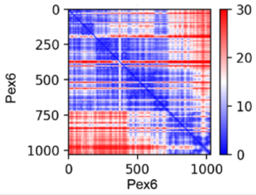  | 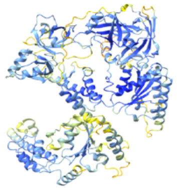 |
